# The structure of arginyltransferase 1 (ATE1)

**DOI:** 10.1101/2022.07.20.500667

**Authors:** Verna Van, Nna-Emeka Ejimogu, Toan S. Bui, Aaron T. Smith

## Abstract

Eukaryotic post-translational arginylation, mediated by the family of enzymes known as the arginyltransferases (ATE1s), is an important post-translational modification that can alter protein function and even dictate cellular protein half-life. Multiple major biological pathways are linked to the fidelity of this process, including neural and cardiovascular developments, cell division, and even the stress response. Despite this significance, the structural, mechanistic, and regulatory mechanisms that govern ATE1 function remain enigmatic. To that end, we have used X-ray crystallography to solve the first crystal structure of ATE1 from *Saccharomyces cerevisiae* ATE1 (*Sc*ATE1) to 2.85 Å resolution. The three-dimensional structure of *Sc*ATE1 reveals a bilobed protein containing a GCN5-related N-acetyltransferase (GNAT) fold, and this crystalline behavior is faithfully recapitulated in solution based on size-exclusion chromatography-coupled small angle X-ray scattering (SEC-SAXS) analyses and cryo-EM 2D class averaging. Structural superpositions and electrostatic analyses indicate this domain as the location of catalytic activity and tRNA binding, and these comparisons strongly suggest a mechanism for post-translational arginylation. Additionally, our structure reveals the spatial connectivity of the N-terminal domain, which we have previously shown to bind a regulatory [Fe-S] cluster, and the enzymatic active site, hinting at the atomic-level details of the cluster’s regulatory influence. When taken together, these insights into the first structure of ATE1 bring us closer to answering pressing questions regarding the molecular-level mechanism of eukaryotic post-translational arginylation.

## INTRODUCTION

Post-translational modifications (PTMs) are important covalent modifications of proteins that can target nearly 75% of amino acid side chains and are essential in affording the expansion of the encoding capacity of the genome.^1^ While some chemical modifications of nascent polypeptides are observed in prokaryotes, PTMs are more frequent in eukaryotes, as evidenced by the significant portions of eukaryotic genomes (*ca*. 5%) dedicated to carrying out these processes.^1^ Several notable PTMs are utilized to control eukaryotic cellular function, including acetylation^2-4^ and methylation^5^ (regulating epigenetics), glycosylation^6^ (regulating cellular signaling and communication), and phosphorylation^7^ (regulating protein oligomerization and signal cascades). These chemical modifications have been heavily studied over multiple decades, and while further research into these essential processes is undoubtedly necessary, it can be argued that these PTMs are well-understood.^1^ However, these examples represent only a fraction of chemical modifications that occur to polypeptides within the cell.^8^

An understudied post-translational modification of emerging importance in eukaryotes is that of arginylation, which is the tRNA-dependent, post-translational addition of the amino acid arginine (Arg) to a polypeptide. This process was first discovered to occur in a non-ribosomal manner in the early 1960s,^9-10^ after which it was shown that this PTM was enzymatically catalyzed^11^. Later, the gene responsible was cloned from *Saccharomyces cerevisiae*, and the encoded enzyme was named arginyltransferase 1, or ATE1^12^. It is now known that this gene is conserved among nearly all eukaryotes, and ATE1-catalyzed post-translational arginylation has begun to emerge as a major player in eukaryotic homeostasis through two different, major functions^13^: its participation in the N-degron pathway, and its role in non-degradative arginylation (Fig. 1). In a degradative manner, ATE1 plays a major role in the Arg branch of the N-degron pathway, a hierarchy that links the identity of the N-terminal residue to its *in vivo* half-life.^14^ In short, ATE1 recognizes proteins bearing negatively-charged, secondary destabilizing residues (Asp, Glu, and Cys-sulfinic acid) on their N-termini and catalyzes N-terminal arginylation in an Arg tRNA^Arg^-dependent manner.^13, 15-16^ These arginylated proteins can be then identified by E3 ubiquitin ligases and tagged for degradation via the proteosome.^14^ Essential cellular processes regulated by this pathway include chromosomal segregation^17^, the stress response in yeast^18^, and even cardiovascular development^19^ (Fig. 1). In a non-degradative manner, ATE1 plays a major role whereby a protein is post-translationally arginylated, but this modification stabilizes the polypeptide and can even change its oligomerization propensity, especially in mammalian cells.^20^ Notable examples of ATE1-catalyzed non-degradative arginylation include its established roles in the stabilization of α-synuclein^21^ and β-amyloid^22^, as well as the requirement of arginylation for polymerization of β-actin^23-24^ (Fig. 1). Moreover, it is known that ATE1 is linked to the ability of plants to sense O_2_ levels and to regulate their development in an ATE1-dependent manner (Fig. 1).^25-27^ Thus, a clear importance for ATE1-catalyzed arginylation has been established at the cellular level for virtually all eukaryotes.

**Figure 1.**
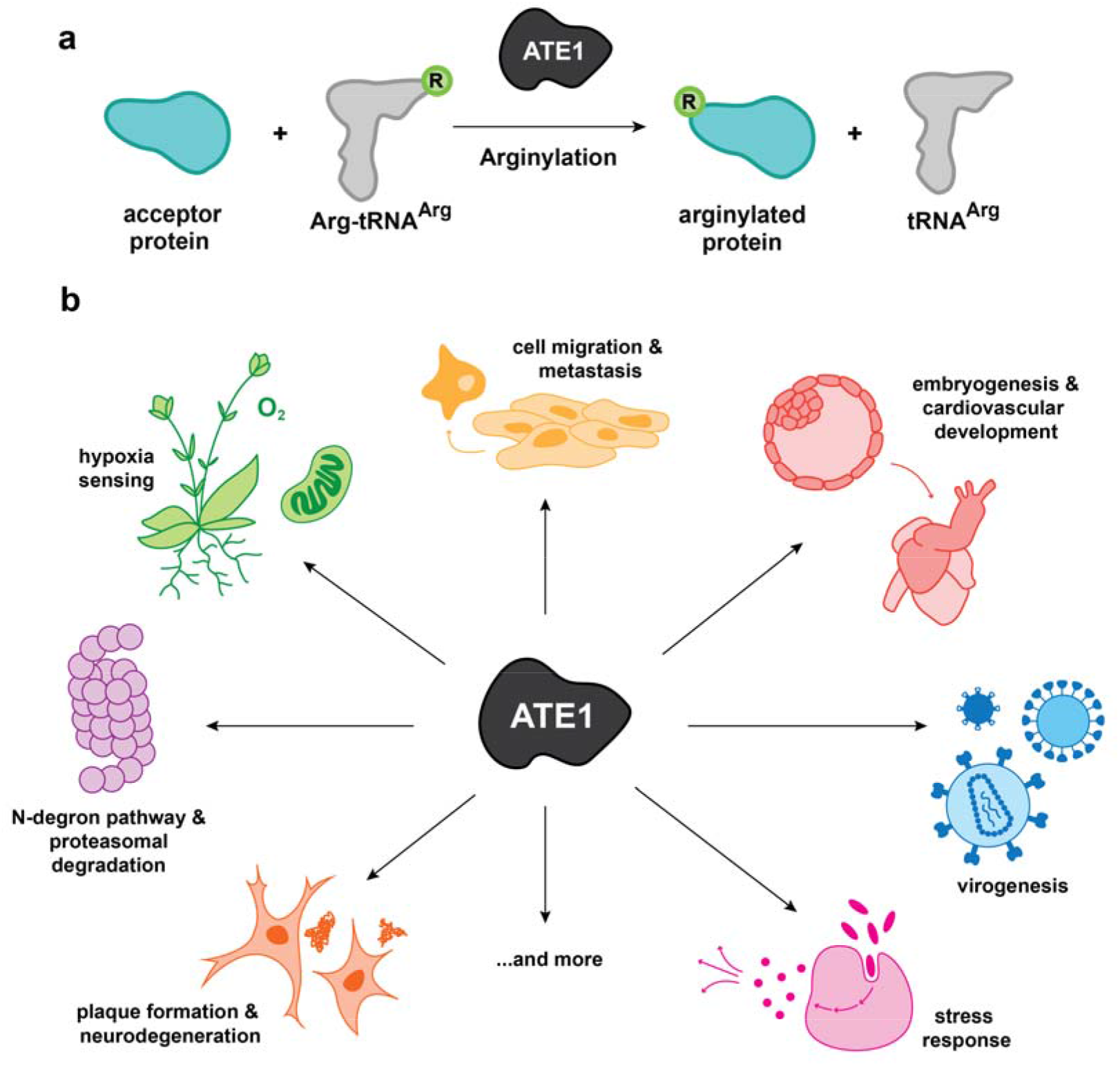
Post-translational arginylation and its impacts. **a**. Cartoon depiction of the arginylation reaction. **b**. ATE1-catalyzed post-translational arginylation is a key regulator of eukaryotic homeostasis. ATE1 catalyzes post-translational arginylation, which is the covalent addition of the amino acid Arg to a polypeptide bearing acidic N-terminal residues such as Asp, Glu, or Cys-sulfinic acid, although reports indicate that arginylation can also occur at certain points on the interior of the polypeptide (mid-chain arginylation). The arginylation of a protein regulates its fate and its function in degradative and non-degradative ways. Examples of key cellular processes that depend on protein arginylation include: cell migration and metastasis, embryogenesis and cardiovascular development, virogenesis, the stress response, plaque formation and neurodegeneration, the N-degron pathway and proteasomal degradation, and even hypoxia sensing.

Despite this significance, major structural, mechanistic, and regulatory questions regarding ATE1 and N-terminal arginylation remain outstanding. Notably, while attempts have been made to understand the ATE1 enzymatic active site^28-30^ and its domain topology^13^, and even to determine models of ATE1 based on loose sequence homology^31^, the structure of ATE1 has remained undetermined. Moreover, these low-resolution techniques and the lack of a structure have hampered the generation of a consensus mechanism of N-terminal arginylation. An additional complication to the determination of ATE1 structure is our recent discovery that ATE1s are O_2_-sensitive, [Fe-S] cluster-binding enzymes.^32^ While we mapped the binding of this [Fe-S] cluster to the N-terminal regulatory domain of ATE1, demonstrated its effects on *in vitro* arginylation, and showed its evolutionary conservation, we noted that heterologous expression of ATE1 in *Escherichia coli* could result in heterogeneous loading of the [Fe-S] cluster if not precisely controlled.^32^ This heterogeneity could hamper the formation of sufficient homogeneous protein for structural characterization. Additionally, as ATE1 itself can be prone to aggregation, reacts with multiple protein substrates, and uses Arg tRNA^Arg^ as a co-substrate,^13^ it is perhaps unsurprising that structural insight into this enzyme has been slow.

In this work, we report the first three-dimensional structure of apo ATE1 from *Saccharomyces cerevisiae* (*Sc*ATE1) determined to 2.85 Å resolution. The X-ray crystal structure reveals that *Sc*ATE1 is a bilobed protein, and this overall topology is confirmed in solution by using small-angle X-ray scattering (SAXS) and cryo-EM 2D class averaging. Surprisingly, the smaller lobe of *Sc*ATE1 is the location of the GCN5-related *N*-acetyltransferase (GNAT) fold that houses the enzymatic active site, while the larger lobe is composed of the N-terminal regulatory domain (site of [Fe-S] cluster binding) folded against the C-terminal domain of unknown function. Based on structural superpositions with prokaryotic functional analogs (L/F-transferases) and electrostatic analyses, we posit that the GNAT fold binds the 3’ arginylated stem of Arg tRNA^Arg^ in a polar cleft terminating at a locus of negative charge (Asp^277^), which is highly conserved in this location. Based on conservation of residues and on comparisons to the L/F-transferase mechanism, we hypothesize the first mechanism of ATE1-catalyzed N-terminal arginylation derived from atomic-level structural information. Taken together, these results provide the first structure of an ATE1 and lend insight into the mechanism of this important PTM, which could be leveraged as a target for future therapeutic interventions.

## RESULTS

### Crystallization and Phasing

The phasing of purified and crystallized apo *Sc*ATE1 (Supplementary Figure 1) was a particularly noteworthy endeavor. Both intact and tag-cleaved *Sc*ATE1 crystallized within space group *P*4_3_ composed of a large unit cell (a = b = 235.27 Å; c = 171.10 Å; α = β = γ = 90°), with the best native crystals exhibiting diffraction resolved to 2.85 Å (Supplementary Table 1). A Matthew’s analysis of the unit cell indicated incredibly broad and ambiguous solvent and asymmetric unit (ASU) contents, with the putative number of polypeptides in the ASU ranging from 8-20 molecules (*ca*. 75-40 % solvent content). Initial attempts to phase the data using molecular replacement (MR) of prokaryotic functional analogs known as L/F-transferases (*e*.*g*., PDB ID 2CXZ and 2Z3K) were unsuccessful, likely owing to the poor sequence conservation between these enzymes and ATE1s (*ca*. 10% sequence identity; < 20% sequence similarity). Multi-wavelength anomalous dispersion (MAD) of Pt-, Au-, or Hg-metallated derivatives of crystallized, native ATE1 failed to diffract isotropically to resolutions better than 6 Å. Phasing was then attempted via single-wavelength anomalous dispersion (SAD) using SeMet-derivatized ATE1. Despite good accumulation and purity, crystals of SeMet ATE1 were small and still diffracted poorly (*ca*. 4 Å resolution at best) with marginally resolvable anomalous signal due to rapid radiation damage. Coupled with the ambiguity of the ASU contents and the low anomalous signal, phasing initially seemed intractable. However, the recent release of the AlphaFold EMBL database^33^ included a predicted high confidence model of full-length *Sc*ATE1, which we then used for MR testing. After model truncation to remove low-confidence regions, we applied a bootstrapping approach to MR wherein we started with the placement of a single *Sc*ATE1 protomer model and continued to increase the ensemble size based on the increasing Phaser score and the presence of clear, positive *F*_o_-*F*_c_ difference density in the map of the ASU. This approach converged to a single MR result (TFZ score of 29.6; LLG score of 5472) with 12 molecules placed in the ASU, corresponding to a reasonable solvent content of 63.5%. Attempts to increase the composition further failed, as evidenced by drastically lowered TFZ and LLG scores, large amounts of clear, negative *F*_o_-*F*_c_ difference density, and unrealistic packing clashes. Thus, our phased dataset with 12 molecules placed in the ASU was used for subsequent rebuilding and refinement, eventually resulting in a final model exhibiting *R*_w_ and *R*_f_ values of 0.199 and 0.257, respectively (Supplementary Table 1).

### Overall architecture

The unit cell of crystallized *Sc*ATE1 is very large (*ca*. 9.5 million Å^3^), and the ASU is composed of 12 arginine transferase protomers (Supplementary Figure 2), each of which adopts a bilobed structure (Fig. 2). Within the ASU of apo *Sc*ATE1 resolved to 2.85 Å, 5493 amino acids, 203 SO_4_^2-^ anions, 41 glycerol molecules, and 64 H_2_O molecules were all visible in the electron density (Supplementary Figure 3). Each *Sc*ATE1 protomer is composed of 22 α helices/helical turns and 19 β strands with the approximate dimensions of ∼70 Å by ∼40 Å by ∼40 Å without any solvent considerations, and a surface view of the protein roughly resembles the shape of the country of Australia (Fig. 2a-d). In general, the α helices pack the outside of the protein, with three mutually perpendicular β sheets running through the central, longest axis of the polypeptide. There are four dynamic regions within the crystal that were unable to be modeled due to the lack of electron density: residues 17-31 (part of the N-terminal regulatory domain); residues 295-306 (part of a predicted helical turn connecting β11 and β12); 413-421 (a surface-exposed predicted turn that is part of the C-terminal domain directly abutting the N-terminal regulatory domain); and 503-518 (a TEV cleavage site and the His_6_ tag, disordered within a solvent channel) (Fig. 2a,b). There are minimal variations (± 1-2 amino acids visible in the electron density) of these disordered regions amongst the 12 protomers, and superpositioning indicates high structural similarity: the most structurally-similar protomers exhibit < 0.5 Å RMSD across all C_α_s, and the most structurally-disparate protomers exhibit *ca*. 0.7 Å RMSD across all C_α_s. Based on fit to the electron density and the lowest overall thermal factors, the highest-quality protomers within the structure appear to be those of chains A, C, and E (Supplementary Figures 2 and 3). Despite the large ASU, the placement of the protomers does not appear to be due to any obvious non-crystallographic symmetry or any conserved quaternary structure; rather, the crystal packing appears to be random in the ASU, which agrees with data demonstrating the *Sc*ATE1 behaves monomerically in solution (*vide infra*).

**Figure 2.**
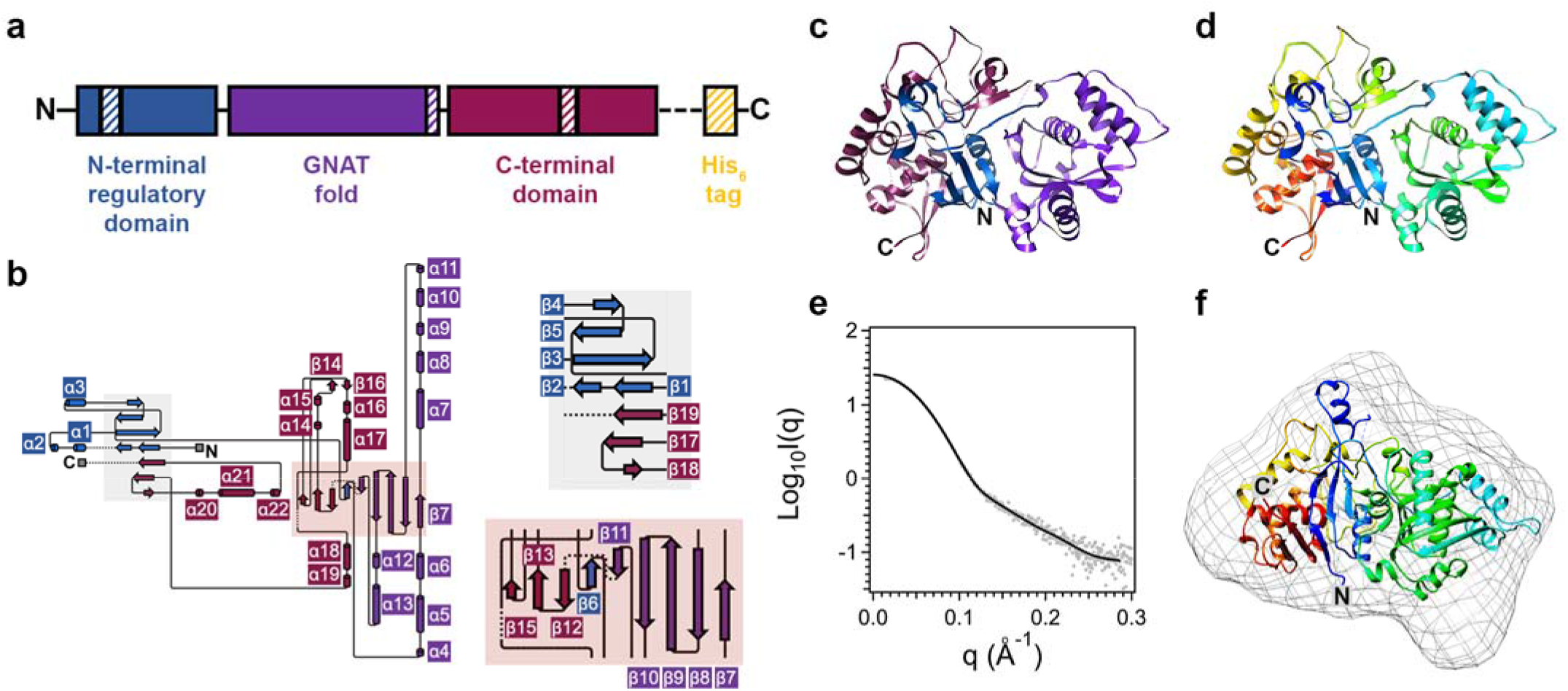
The structure of *Saccharomyces cerevisiae* ATE1 (*Sc*ATE1). **a**. Schematic cartoon of the *Sc*ATE1 domain organization. The N-terminal regulatory domain is colored blue, the GNAT fold is colored purple, the C-terminal domain (unknown function) is colored magenta, and the cleavage site and His_6_ tag are colored yellow. Hashed rectangles represent locations of disorder in the structure. **b**. Secondary-structure topology diagram of a single *Sc*ATE1 polypeptide. α helices and β sheets are numbered sequentially. The β sheet regions are enlarged and placed in boxes colored gray and pink for clarity. **c**. The 2.85 Å X-ray crystal structure of a single apo *Sc*ATE1 protomer, color-coded the same as panel (**a**). **d**. The 2.85 Å X-ray crystal structure of a single apo *Sc*ATE1 protomer, color-coded blue (N-terminus) to red (C-terminus). **e**. The log_10_ plot of apo *Sc*ATE1 SEC-SAXS data indicating that the protein is monodisperse with negligible aggregation. **f**. Overlay of a single apo *Sc*ATE1 protomer with the *ab initio* envelope (gray mesh) generated from the SEC-SAXS data. In all cases ‘N’ and ‘C’ represent labels for the N- and C-termini, respectively.

The overall shape of apo *Sc*ATE1 is confirmed by small-angle X-ray scattering analyses (SAXS) and cryo-EM 2-D class averaging. To determine the overall shape and monodispersity of *Sc*ATE1 in solution, we performed multi-angle light scattering (MALS) and size-exclusion-coupled SAXS (SEC-SAXS). The experimental SAXS data (Fig. 2e) indicate good monodispersity in solution, and the Kratky plot for apo *Sc*ATE1 indicates a roughly globular conformation (Supplementary Figure 4). MALS data confirm our gel filtration experiments, showing that apo *Sc*ATE1 behaves monomeric in solution. We were then able to generate an *ab initio* envelope based on these high-quality data (Fig. 2f), and an overlay of a single *Sc*ATE1 protomer fits nicely within the envelope. While our SAXS data suggest that there is some dynamicism in a C-terminal loop that is pinned against the body of the protein in the X-ray structure (but may be free or at least more dynamic in solution), there is a general and faithful recapitulation of the overall shape of the *Sc*ATE1 protomer in solution compared to the X-ray structure. To further confirm these observations, cryo-EM grids of *Sc*ATE1 were frozen and images were initially analyzed in collaboration with the Hauptman-Woodward Institute (HWI)’s Cryo-EM center. Data processing proceeded through particle picking, and after several rounds of 2D classifications, 13 class averages containing 78,153 particle images were selected for further refinement. Although 3D reconstruction and map refinement of *Sc*ATE1 to a resolution better than 4 Å was ultimately unsuccessful, it is still evident that the resultant 2D class averages show particle images that strongly resemble the bilobed structure ascertained via SAXS and X-ray crystallography (Supplementary Figure 5). Thus, in combination with our SAXS data our cryo-EM 2D classes averages, there appears to be excellent agreement between the solution-state and crystalline behavior of *Sc*ATE1.

### Active and regulatory sites

The crystal structure of *Sc*ATE1 reveals the presence of the transferase enzymatic active site, which is housed surprisingly within the smaller protein lobe of ATE1 (Figs. 2a-c, 3a). The presence of a GCN5-related *N*-acetyltransferase (GNAT) fold in the crystal structure of apo *Sc*ATE1 is unmistakable and is roughly composed of residues 107-307 (Figs. 2b, 3a). The lobe containing the GNAT fold comprises 10 α helices/helical turns (α4−α13) and 5 β strands (β7−β11) resembling an α/β/α sandwich (Fig. 3a). While a typical GNAT fold has 5 β strands comprising an antiparallel β sheet similar to the *Sc*ATE1 active site (Fig. 3a), most GNAT folds have only 3-4 α helices/helical turns, far fewer than those observed in *Sc*ATE1.^34^ The antiparallel β sheet made of β strands 7-10 are “sandwiched” between α7/α8 on one face, while the opposite face is capped by α5 and α6 (connecting to the N-terminal domain) as well as α13; the remaining α helices/helical turns flank the sides of the fold (Figs. 2b, 3a). Two β strands β10 and β11 of the GNAT fold run parallel and open apart to form a V-shaped cleft as they exit the α/β/α sandwich (Figs. 2b, 3a), which is a common characteristic of the substrate(donor)-binding site of the GNAT fold. Indeed, superpositioning of *Sc*ATE1 with the *Escherichia coli* L/F-transferase complexed with phenylalanyl adenosine^35^ (rA-Phe, a proxy for the 3’ end of phenylalanylated tRNA; PDB ID 2Z3K; Fig. 3c) reveals a general conservation of the GNAT fold between the two functional analogs (*ca*. 3.2 Å RMSD over 188 C_α_s; Fig. 3c) and points to these two strands that are splayed apart as the location for amino acid binding of the 3’ end of the aminoacylated tRNA (Figs. 3d,e). Wholly consistent with their differences in which aminoacylated tRNA(s) is/are sensed as donor substrates, the Phe sidechain of rA-Phe binds within a modest hydrophobic pocket (*ca*. 5 Å in depth) in the *E. coli* L/F-transferase (Fig. 3d), while *Sc*ATE1 has a deeper (*ca*. 8 Å), more polar pocket (lined by residues Thr^244^, Ser^246^, Ser^255^, Ser^273^, and Tyr^289^) that terminates at the carboxylate side chain of Asp^277^ (Figs. 3b, 3d). A negative locus of charge in this pocket is likely necessary to recognize the increased size and complementary positive charge of the Arg side chain for ATE1s (Fig. 3f), and this observation is supported by multiple sequence alignments demonstrating conservation of either Asp or Glu in this position (Supplementary Figure 6). Leading into this polar pocket is a bulky, polar gate composed of Tyr^293^ and Tyr^294^, while the side of the active-site pocket is blocked by a hydrophobic flank composed of Phe^258^ and Trp^260^ (Fig. 3a) that likely prohibits entry of substrate at the incorrect angle. Modeling and superpositioning of rA-Arg onto rA-Phe (PDB ID 2Z3K; Fig. 3e) indicates that the aminoacylated tRNA containing the guanidinium sidechain would be within H-boding distance of the Asp^277^ carboxylate (*ca*. 3 Å; Fig. 3f), although the exact conformer that Arg would adopt in this pocket is unknown at this time.

**Figure 3.**
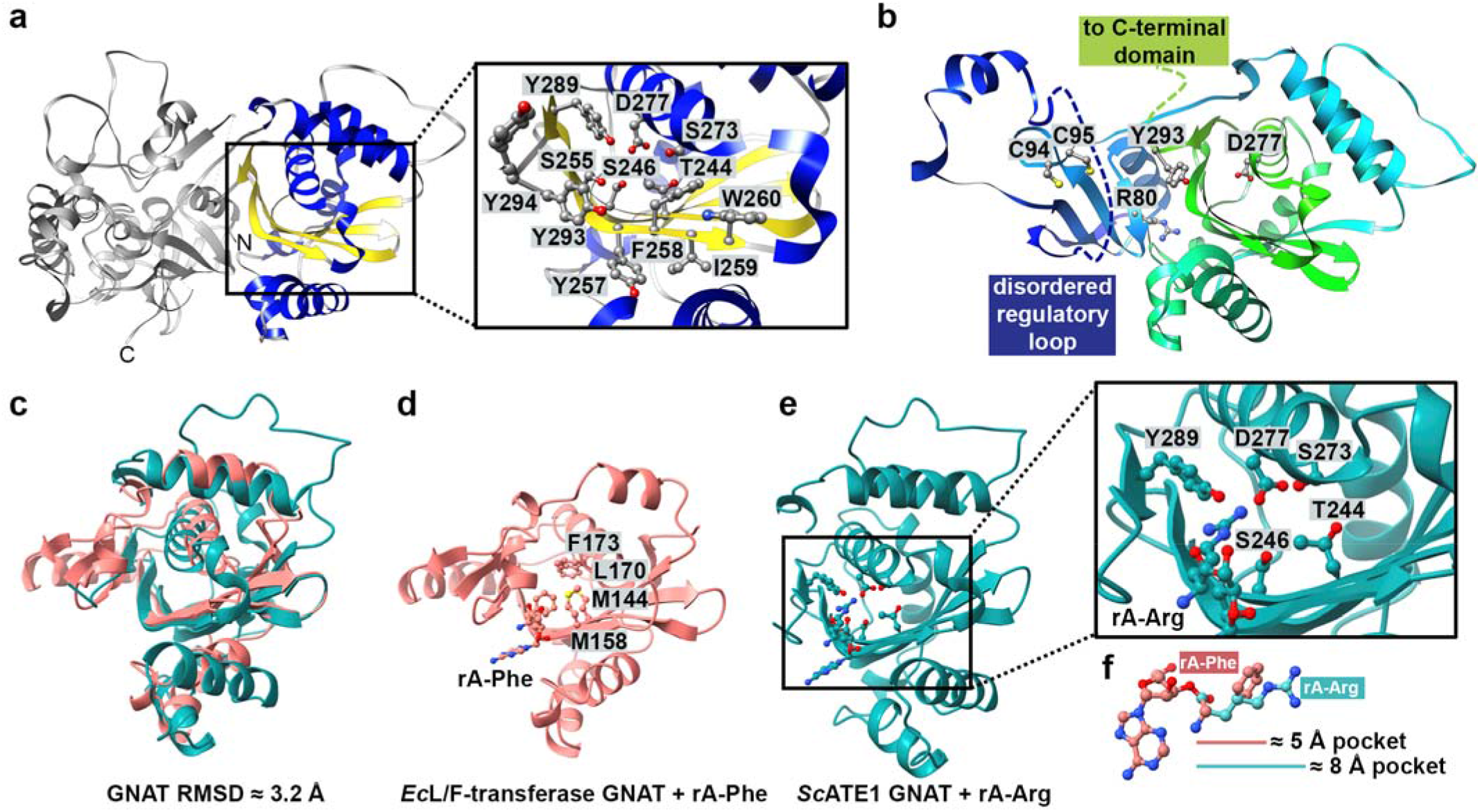
The enzymatic active site of *Sc*ATE1. **a**. A single protomer of *Sc*ATE1 with the GNAT fold highlighted in blue and yellow. Inset: key residues of the polar pocket of the *Sc*ATE1 GNAT fold and the flanking hydrophobic blockade. ‘N’ and ‘C’ represent the N- and C-termini, respectively. **b**. Residues 1-295 of *Sc*ATE1 illustrating: the N-terminal regulatory domain (including Cys^94^ and Cys^95^); the putative residue that engages the substrate protein, Arg^80^; the putative H-bonding partner of the aminoacyl carbonyl, Tyr^293^; and the residue of interaction with the 3’ Arg residue, Asp^277^. **c**. Overlay of the GNAT folds of the *E. coli* L/F-transferase (PDB ID 2Z3K; salmon) and that of *Sc*ATE1 (teal). The RMSD is ≈ 3.2 Å across 188 C_α_s. **d**. The GNAT fold of the *E. coli* L/F-transferase complexed with rA-Phe, which binds in a hydrophobic pocket ≈ 5 Å deep. **e**. The GNAT fold of the *Sc*ATE1 superposed with rA-Arg. Inset: rA-Arg is predicted to bind within the deeper hydrophilic pocket (≈ 8 Å) of the *Sc*ATE1 GNAT fold and make electrostatic and H-bonding interactions with Asp^277^ and Tyr^289^, respectively. **f**. Comparative sizes of rA-Phe (salmon) and rA-Arg (teal) and their respective binding pocket sizes in the *E. coli* L/F-transferase and *Sc*ATE1 GNAT folds.

Intriguingly, the crystal structure of *Sc*ATE1 reveals that the N- and C-terminal domains pack together to form the larger protein lobe (Figs. 2b-d, 3a), which was unexpected. The larger portion of the bilobed *Sc*ATE1 protomer is composed roughly of amino acids 1-103 (the N-terminal regulatory domain) (Figs. 2a, 3b) and amino acids 307-502 (the C-terminal domain of unknown function) (Figs. 2a, 3a). A search of the Dali protein structural comparison server^36^ revealed no obvious structural similarity of either of these portions of ATE1 to other known proteins, and the function of the C-terminal domain remains unclear. However, we were able to visualize the partial location of the [Fe-S] cluster-binding site in the N-terminal domain (Fig. 3b), which we have previously shown to be important for the regulation of arginylation^32^. Based on our previous work, we have demonstrated that the [Fe-S] cluster-binding site is composed of Cys^23^ and Cys^26^ (part of a conserved CGYC motif) and Cys^94^/Cys^95^ (part of a conserved CC motif).^32^ While we have yet been able to crystallize the holo (cluster-replete) form of the protein, the AlphaFold model of *Sc*ATE1 supports co-localization of all four Cys residues in spatial proximity. Our apo structure reveals that the CGYC motif is part of a dynamic and disordered loop stretching from residues 15-33 (Supplementary Figure 7), and we speculate that this region becomes ordered in the presence of the [Fe-S] cluster. We can clearly see the reduced Cys^94^/Cys^95^ vicinal pair, which lies along a region of random coil running directly into the GNAT fold, antiparallel to β11 of the V-shaped cleft within the GNAT active site, and directly connected to helices α5 and α6 that cap the active site (Fig. 3b, Supplementary Figure 7). These observations definitively demonstrate that these Cys residues are not part of the enzymatic active site *per se* (as previously suggested in ^28-29^), but rather link to the active site through critical contacts at the lobe-lobe interface (Fig. 3b). Moreover, it is easy to imagine how changes in the status of the cluster in the N-terminal domain could be transmitted along this interface to change access to the GNAT fold in a subtle but instrumental way. It is tempting to speculate that merbromin (a known ATE1 inhibitor)^37^ could bind in a similar location as the [Fe-S] cluster in the N-terminal domain and affect arginylation activity, although further work is necessary to support this hypothesis.

The surface structure and the uneven charge distribution of apo *Sc*ATE1 reveal a putative site for the binding of a portion of the tRNA macromolecule (Supplementary Figures 8,9). Visualization of the protein surface shows a clear and obvious cleft at the interface between the two protein lobes (Supplementary Figure 8) that funnels into the electronegative patch that terminates at Asp^277^ (Supplementary Figure 9). While the cleft between these two lobes is sizable (*ca*. 6700 Å^3^; Supplementary Figure 8), an intact aminoacylated tRNA would not fit within this volume. However, it is possible that a tRNA fragment could fully bind in this area (as previously suggested)^38^, or that only a portion of the acceptor stem makes contact with this region. Furthermore, the electrostatics of the smaller lobe of the protein and the lobe-lobe interface are generally electropositive (particularly along the surface-exposed regions of α7 and α5; Supplementary Figure 9), whereas the larger C-terminal lobe of the protein is highly electronegative, which would prohibit the binding of the aminoacylated tRNA (either intact or fragmented) due to electrostatic repulsion of RNA phosphate backbone.

Despite our structure providing clear indicators of where the 3’ arginylated tRNA binds to the GNAT fold and where the larger tRNA macromolecule engages the protein, the binding site within *Sc*ATE1 of the carboxylate or sulfinic acid side chain of the secondary destabilizing residue on the substrate polypeptide was less clear. As the arginylated tRNA likely occupies a substantial fraction of the cleft volume at the lobe-lobe interface, it is unlikely that this space could also be occupied by the substrate polypeptide. A structural analysis of other regions of *Sc*ATE1 that might engage the substrate polypeptide resulted in four likely candidates: Arg^79^ and Arg^80^, two vicinal Arg residues located on the opposite face of the V-shaped cleft relative to Asp^277^; His^231^, located along β8 and on the opposite face of the GNAT β sheet as Asp^277^; and Arg^276^, located one residue removed from Asp^277^ and pointing opposite (away) from the cleft at the lobe-lobe interface as Asp^277^. We used multiple sequence alignments to determine which of these positively-charged residues were highly conserved (Supplementary Figure 6), and Arg^79^ and Arg^80^ were the two most likely candidates. Based on our structural data and docking, we believe Arg^80^ is the most likely candidate to engage in electrostatic interactions between *Sc*ATE1 and the negatively-charged side chain on the protein substrate.

### Proposed mechanism and functional implications

Our structural and biochemical data allow us to propose a mechanism of post-translational N-terminal arginylation (Fig. 4). Based on surface electrostatics, we believe that the positively-charged GNAT surface is involved in the initial electrostatic interactions with the phosphate backbone of the aminoacylated tRNA, and at least some of the tRNA may be accommodated within the sizable cleft between the two ATE1 lobes. The 3’ end of Arg-tRNA^Arg^ is forced to assume the correct orientation by a hydrophobic blockade (Phe^258^ and Trp^260^) and is guided into the V-shaped cleft of the GNAT fold by hydrophilic residues lining a polar pocket that terminates in Asp^277^ (highly conserved as either Asp or Glu in this position). On the opposite face of ATE1 is the location of what we posit to be the entry site for the protein substrate bearing a secondary destabilizing residue at its N-terminus (Asp, Glu, or Cys-sulfinic acid). Direct electrostatic engagement with the highly conserved Arg^80^ would lock the negatively-charged side chain in place to position the N-terminal amine to react with the aminoacyl bond. The tetrahedral intermediate with its oxyanion is likely stabilized via H-bonding with Tyr^293^ (or the vicinal Tyr^294^), analogous to the function of Asn^191^ in the *E. coli* L/F-transferase. Collapse of the tetrahedral intermediate and proton transfer would yield a new N-terminus via formation of a canonical peptide bond in a non-ribosomal manner and would free the 3’-OH of the tRNA (Fig. 4). While the driving forces for release of the N-terminally arginylated substrate protein and the free tRNA are currently unknown, this proposed mechanism bears a strong resemblance to that of a similar mechanism proposed by Watanabe *et al*. to explain the general N-terminal peptide-bond formation catalyzed by the prokaryotic L/F-transferases. However, unlike in the L/F-transferases where a proton relay network comprising Gln^188^ and Asp^186^ is proposed to be operative, analogous positions in ATE1 are either hydrophobic or do not point in the proper orientation to conserve this portion of the mechanism in ATE1. In contrast, given the generally more hydrophilic GNAT pocket, it may be that any number of the hydrophilic residues (and their H-bonding network) lining this region may serve a similar purpose, or that the tRNA 3’ ribose hydroxyl might be involved, as has been proposed as an alternative mechanism of the L/F-transferases^39^. Future work will be necessary to decipher whether either (or any) of these possibilities are operative.

**Figure 4.**
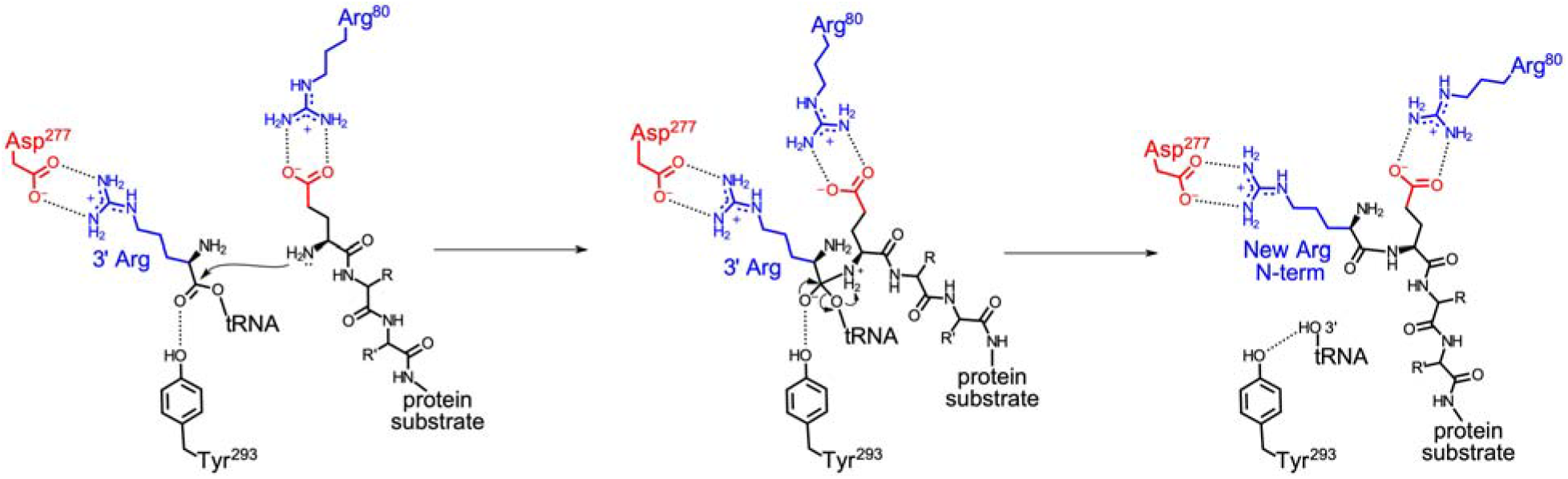
Proposed mechanism of ATE1-catalyzed post-translational N-terminal arginylation, based on our structural and biochemical work. In addition to electrostatic interactions of the tRNA backbone with the electropositive GNAT fold, the 3’ Arg on the aminoacylated tRNA is recognized and bound via electrostatic interactions with Asp^277^ that position the amino-acyl bond for nucleophilic attack of the N-terminus (left panel). Arg^80^ is likely involved in recognition of the negatively-charged side chain of the N-terminal secondary-destabilizing residue (Asp, Glu, or Cys-sulfinic acid), while Tyr^293^ plays an analogous role to Asn^191^ of the L/F-transferases to stabilize the tetrahedral intermediate formed between the aminoacylated tRNA and the protein substrate (middle panel). Collapse of the tetrahedral intermediate and proton transfer liberates the tRNA 3’ hydroxyl (-OH) and the N-terminally arginylated protein substrate (right panel). This process completes the N-terminal peptide bond formation in a non-ribosomal manner.

## DISCUSSION

The importance of post-translational arginylation has become increasingly evident over the past three decades. First discovered as an intriguing method of incorporating Arg into mature proteins in a non-ribosomal manner,^9-10^ ATE1 and tRNA-dependent arginylation have emerged as global regulators of eukaryotic homeostasis (Fig. 1).^13^ One way in which arginylation is essential is via its role in the N-degron pathway, a hierarchical pathway that links the identity of the N-terminus to protein half-life via the ubiquitin-proteasome system.^15-16, 40^ While some controversy surrounds which proteins are *bona fide* targets of the N-degron system, it is clear that this pathway plays an important role in cardiovascular development^19^, neurogenesis^41^, and even G-protein signaling^42^, among other biological processes (Fig. 1). Another way in which arginylation is essential is in its ability to alter and to control protein oligomerization in a non-degradative manner. The most notable example of this type of oligomerization control is β-actin, which is an essential cytoskeletal protein that depends on arginylation for proper function^23^, although several other examples highlighting the importance of non-degradative arginylation also exist^11, 21, 26-27, 43-44^ (Fig. 1). Broadening its importance, recent studies have even demonstrated links between ATE1-catalyzed arginylation and certain types of cancer^45^, while connections to the virulence of serious human infectious agents such as the human immunodeficiency virus (HIV)^46^ and severe acute respiratory syndrome coronavirus 2 (SARS-CoV-2; cause of the COVID-19 pandemic) are emerging^47^ (Fig. 1). Despite this significance and focused efforts of several research groups to understand arginylation at the cellular level, information regarding the structure of ATE1, its mechanism of action, and its regulation remained obscure.

In this work, we have determined the first structure of an ATE1—the eukaryotic enzyme that catalyzes post-translational arginylation—and we have proposed its mechanism of action in N-terminal arginylation. The 2.85 Å X-ray crystal structure of ATE1 from *Saccharomyces cerevisiae*, which faithfully recapitulates in-solution behavior as determined by SEC- and MALS-coupled SAXS, revealed a bilobed structure composed of two domains. Housed within the larger lobe of the protein is the N-terminal regulatory domain, which we have previously shown to bind a regulatory [Fe-S] cluster^32^, folded against a C-terminal domain of unknown function, which was unexpected (Fig. 2). Contained within the smaller lobe of the protein is the enzymatic active site, which is composed of a GNAT fold lined with polar residues that funnel the 3’ arginylated end of the Arg tRNA^Arg^ into a cleft that terminates at Asp^277^. Based on superpositioning of the *E. coli* L/F-transferase structure complexed with rAF (PDB ID 2Z3K)^35^, we were able to model the location of the 3’ end of Arg tRNA^Arg^ into this polar pocket, and sequence alignments allowed us to posit the binding model of the substrate protein with that of ATE1 (Fig. 3). Based on these observations, we propose a mechanism in which electrostatic interactions in the GNAT fold (in particular Asp^277^) and on the adjacent N-terminal domain (in particular Arg^80^) serve essential roles in forming a ternary complex that is competent for N-terminal arginylation (Fig. 4). These electrostatic interactions would position the N-terminus of the protein substrate within proximity of the aminoacylated 3’ end, allowing for reaction and amino acid transfer from the charged tRNA in a non-ribosomal manner (Fig. 4). Fascinatingly, this ATE1-catalyzed mechanistic hypothesis bears a striking resemblance to that of the proposed mechanism of the L/F-transferases despite the differences in tRNA and protein substrates that are recognized, and notwithstanding the low sequence similarity between this eukaryotic enzyme and its prokaryotic functional analog.

The determination of the first atomic-level structure of an ATE1 has now set the stage for a greater understanding of ATE1 mechanism and regulation. For example, our recent results have shown that ATE1 is multifaceted by also being an [Fe-S] cluster-binding enzyme. We posited that the presence of this cluster appears to function as an O_2_-dependent regulator to control post-translational arginylation in response to oxidative stress^32^, which is an important aspect of ATE1 function. While our new structure is apo (lacks the [Fe-S] cluster), it is tempting to speculate the mechanistic connection between cluster-binding in the N-terminal regulatory domain and alterations that could occur at the GNAT fold, and a metal-bound structure would significantly increase our understanding of this regulatory paradigm. Additionally, it is known that ATE1 can catalyze mid-chain arginylation, at least under certain cellular contexts.^30^ Based on our current mechanistic hypothesis and structural data, it is currently unclear how this process is chemically possible, unless mid-chain protein substrates access the enzymatic active site in an alternative manner. Finally, a detailed understanding of how ATE1 engages partner macromolecules, such as tRNA or tRNA fragments^38^, or its putative protein binding partner in higher-order eukaryotes (the ligand of ATE1, or Liat1^48^), remains to be uncovered. Structures involving these partner molecules are of great interest, and atomic-level understandings of these macromolecular interactions would reveal a deeper understanding of ATE1 function in the context of more complex components of the cellular milieu.

## Abbreviations

ATE1: arginyltransferase 1
ASU: asymmetric unit
ATP: adenosine triphosphate
GNAT: GCN5-related *N*-acetyltransferase fold
L/F-transferase: leucyl/phenylalanyl-tRNA-protein transferase
MAD: multi-wavelength anomalous dispersion
MR: molecular replacement
PTM: post-translational modification
RMSD: root-mean-square deviation
SAD: single-wavelength anomalous dispersion
SAXS: small-angle X-ray scattering
SEC: size-exclusion chromatography
tRNA: transfer ribonucleic acid

## Acknowledgements

This work was supported by NIH-NIGMS grant R35 GM133497 (A.T.S) and NIH-NIGMS grant T34 GM136497 (N.-E.E.). This research used resources of the Advanced Photon Source, a U.S. Department of Energy (DOE) Office of Science User Facility operated for the DOE Office of Science by Argonne National Laboratory under Contract No. DE-AC02-06CH11357. Use of the LS-CAT Sector 21 was supported by the Michigan Economic Development Corporation and the Michigan Technology Tri-Corridor (Grant 085P1000817). SAXS experiments were conducted at the Advanced Light Source (ALS), a national user facility operated by Lawrence Berkeley National Laboratory on behalf of the Department of Energy, Office of Basic Energy Sciences, through the Integrated Diffraction Analysis Technologies (IDAT) program, supported by the DOE Office of Biological and Environmental Research. Additional support comes from the National Institute of Health project ALS-ENABLE (P30 GM124169) and a High-End Instrumentation Grant S10OD018483. Sequence searches utilized both database and analysis functions of the Universal Protein Resource (UniProt) Knowledgebase and Reference Clusters (http://www.uniprot.org) and the National Center for Biotechnology Information (http://www.ncbi.nlm.nih.gov/).

## Author contributions

V.V., N.-E.E., T.S.B., and A.T.S. designed the research; V.V., N.-E.E., T.S.B. and A.T.S. performed the research; V.V., N.-E.E., and A.T.S. analyzed the data; and V.V., N.-E.E., and A.T.S. wrote and edited the paper.

## Additional information

### Supplementary information

Supplementary information accompanies this paper.

### Competing interests

The authors declare no competing interests.

## METHODS

### Materials

All materials used for buffer preparation, protein expression, and protein purification were purchased from commercial vendors and were used as received.

### Cloning, expression, and purification of *Sc*ATE1 constructs

The cloning, expression, and purification of *Sc*ATE1 followed similar protocols as described in ^32^. Briefly, DNA encoding for the gene corresponding to the single-isoform arginine transferase (ATE1) from *Saccharomyces cerevisiae* (strain ATCC 204508; Uniprot identifier P16639) (*Sc*ATE1) with an additionally engineered DNA sequence encoding a C-terminal TEV-protease cleavage site (ENLYFQS) was subcloned into the pET-21a(+) expression plasmid, encoding a C-terminal (His)_6_ affinity tag when read in-frame. The complete expression plasmid was transformed into chemically competent BL21(DE3) cells, spread onto Luria-Bertani (LB) agar plates supplemented with 100 μg/mL ampicillin, and grown overnight at 37 °C. Expression of *Sc*ATE1 was accomplished in 12 baffled flasks each containing 1 L sterile LB supplemented with 100 μg/mL (final) ampicillin and inoculated with a pre-culture derived from a single bacterial colony. Cells were grown with shaking until the optical density at 600 nm (OD_600_) reached ≈ 0.6 to 0.8. The flasks containing cells and media were then chilled to 4 °C for 2 h, after which protein expression was induced by the addition of isopropyl β-D-l-thiogalactopyranoside (IPTG) to a final concentration of 1 mM. The temperature of the incubator shaker was lowered to 18 °C with continued shaking. After *ca*. 18 h to 20 h, cells were harvested by centrifugation at 4800×*g*, 10 min, 4 °C. Cell pellets were subsequently resuspended in resuspension buffer (50 mM Tris, pH 7.5, 100 mM NaCl, 0.7 mM glycerol), flash-frozen on N_2(l)_, and stored at -80 °C until further use. For the expression of SeMet-substituted *Sc*ATE1, the same protocol as outlined in Ref. ^49^ was followed. Purification of SeMet-substituted *Sc*ATE1 was the same as the native protein (*vide infra*).

All steps for the purification of *Sc*ATE1 were performed at 4 °C unless otherwise noted. Briefly, frozen cells were thawed and stirred until the solution was homogeneous. Solid phenylmethylsulfonyl fluoride (PMSF; 50 mg to 100 mg), and a solution of tris(2-carboxyethyl)phosphine hydrochloride (TCEP; 1 mM final concentration) were added immediately prior to cellular disruption using an ultrasonic cell disruptor. Cellular debris was cleared by ultracentrifugation at 163000×*g* for 1 h. The supernatant was then applied to a metal affinity column that had been charged with Ni^2+^ and equilibrated with 8 column volumes (CVs) of wash buffer (50 mM Tris, pH 8.0, 300 mM NaCl, 1 mM TCEP, and 10 % (v/v) glycerol) with an additional 21 mM imidazole. After washing, protein was then eluted with wash buffer containing 300 mM imidazole. Fractions were concentrated using a 15 mL, 30 kDa molecular-weight cutoff (MWCO) spin concentrator. Protein was then applied to a 120 mL gel filtration column that had been pre-equilibrated with 50 mM Tris, pH 7.5, 100 mM KCl, 1 mM dithiothreitol (DTT), and 5 % (v/v) glycerol. The eluted fractions of the protein corresponding to the protein monomer were pooled and concentrated with a 4 mL, 30 kDa MWCO spin concentrator. Protein concentration was determined using the Lowry assay, and purity was assessed via SDS-PAGE analysis.

### Crystallization and structural determination of *Sc*ATE1

Crystals of native and SeMet-substituted *Sc*ATE1 were obtained by sitting-drop vapor-diffusion using MiTeGen-XtalQuest Plates with a 1:1 (v:v) *Sc*ATE1 (≈ 10 mg/mL) and reservoir solution mixture at room temperature. Medium, three-dimensional, colorless cubes and rods appeared within 24-72 hr and reached their maximal size within 3-5 days on average. Crystals were transferred into cryoprotectant, were soaked for *ca*. 1 min, looped, flash-frozen, and stored at 77 K.

Data sets were collected on beamline LS-CAT 21-ID-F at the Advanced Photon Source, Argonne National Laboratory, using a Rayonix MX-300 CCD detector. Data were processed automatically with Xia2.^50^ The qualities of phases determined using datasets from SeMet *Sc*ATE1 and metal-soaked *Sc*ATE1 were poor and the solvent content was ambiguous, prohibiting the determination of phases using anomalous dispersion. Instead, phases were determined via molecular replacement using Phaser in Phenix^51^ with a truncated initial input of the *Sc*ATE1 AlphaFold model (Uniprot ID P16639). After multiple rounds of bootstrapping with Phaser^51^, extended model building and refinement cycles were performed in Coot^52^ and Phenix Refine^51^, respectively. Final validations were performed using Phenix Validate^51^. The final model consists of 12 polypeptides in the asymmetric unit (ASU), each containing four dynamic regions that were not observed in the electron density and were unable to be modeled: residues 17-31, residues 295-306, residues 413-421, and residues 503-518 comprising the TEV cleavage site and the (His)_6_ tag disordered within a solvent channel. Data collection and refinement statistics are presented in Supplementary Table 1. Structural overlays, Cα RMSD values, and surface representations were generated and calculated by utilizing UCSF Chimera^53^, whereas electron density map overlays were generated using PyMOL. Topology images were generated in part by the Pro-Origami server.^54^ The atomic coordinates for native *Sc*ATE1 have been deposited in the Protein Data Bank (deposition ID 7TIF).

### Small-angle X-ray scattering and cryo-EM imaging of *Sc*ATE1

SEC-coupled small-angle X-ray scattering (SEC-SAXS) data were collected at the Advanced Light Source (ALS), Lawrence Berkeley National Laboratory, on the SIBYLS beamline 12.3.1. A suite of samples each containing 60 µL of *Sc*ATE1 at concentrations ranging from 2-8 mg/mL were screened after passage along a PROTEIN KW-803 column equilibrated with SAXS buffer (50 mM Tris, pH 7.5, 100 mM KCl, 2% (v/v) glycerol, 1 mM DTT) using an autosampler. Eluent was split 2:1 between the X-ray synchrotron radiation source (SAXS) and a series of four inline analytical instruments: 1) Agilent 1260 series multiple wavelength detector (MWD); 2) Wyatt Dawn Helos multi-angel light scattering (MALS) detector; 3) Wyatt DynaPro Titan quasi-elastic light scatterings (QELS) detector; and 4) Wyatt Optilab rEX refractometer. Samples were examined with λ = 1.03 Å incident light at a sample-to-detector distance of 1.5 m resulting in scattering vectors, q, ranging from 0.01 Å^-1^ to 0.5 Å^-1^ where the scattering vector is defined as q=4πsinθ/λ and 2θ is the measured scattering angle. Data were collected in 3 s exposures over the course of 40 min. SEC-SAXS chromatograms were generated and initial SAXS curves were analyzed using SCÅTTER ^55-56^. Additionally, UV, MALS, QELS, and differential refractive index data were collected and analyzed. Scattering curves were analyzed using SCÅTTER^55-56^ and Primus^57^ to generate Guinier and Kratky plots and to determine the radius of gyration (*R*_*g*_) and the maximum particle dimension (*D*_*max*_). *Ab initio* molecular envelopes were generated using DAMMIN^58^ and averaged with DAMAVER^59^ from the ATSAS package and were displayed using UCSF Chimera. *Sc*ATE1 protomers were overlayed with SUPCOMB^60^, also part of the ATSAS package and displayed using UCSF Chimera.

Cryo-EM sample preparation, grid vitrification, grid screening, data collection, data reduction, and 2D-class averaging were all accomplished in collaboration with the mail-in cryo-EM center available in collaboration with the Hauptman-Woodward Medical Research Institute (HWI) based out of the University of Buffalo. Briefly, stock *Sc*ATE1 was prepared at 5 mg/mL in 50 mM Tris, pH 7.5, 100 mM KCl, 1 mM DTT, and 1 % (w/v) trehalose. After dilution to *ca*. 1-2 mg/mL, the protein was deposited in a cold chamber at >90 % humidity onto Cu grids supporting a carbon film, blotted, and vitrified using a Thermo Fisher Vitrobot. Cryo-EM grids were screened and data were collected using a Thermo Fisher Glacios 200 kV microscope with a Volta phase plate equipped with a Falcon 4 Detector. Cryo-grids were imaged at 190,000× magnification (resulting in a pixel size of 0.526 Å on the sample scale) with a defocus range of - 0.9 μm to -2 μm. A total accumulated dose of 56.2 electrons per Å^2^ was exposed across 31 frames and a total of 2.85 s. Acquired movies were reference and gain corrected. The Contrast Transfer Function (CTF) was estimated and corrected using CTFFIND4. After several rounds of 2D classifications using a masking radius of 60 Å, 13 class averages containing 78,153 particle images were selected for further refinement. The 78,153 images were used for further image processing in the SPARX and RELION EM image processing packages. Initial reference maps were generated using cisTEM; however, despite multiple attempts at low-pass filtering and further refinement, the map refinement failed to refine to a resolution better than 4 Å.

### Bioinformatics

All ATE1 sequences were obtained from the Universal Protein Resource (UniProt) Knowledgebase and Reference Clusters (http://www.unprot.org) or the National Center for Biotechnology Information (http://www.ncbi.nlm.nih.gov/). Sequences were retrieved by standard protein-protein BLAST searches (blastp) using *Sc*ATE1 as an input. Conserved amino acids were identified via multiple sequence alignments that were performed using JalView v. 2.7 ^61^ implementing the ClustalW algorithm and the Blosum62 matrix ^62-63^. Sequence logos were generated using Weblogo 3.^*64-65*^

### Data availability

Data are available from the corresponding author upon reasonable request.

## Notes

### Competing Interest Statement

The authors have declared no competing interest.

## References

1. Walsh, C. T.; Garneau-Tsodikova, S.; Gatto, J., Gregory J., Protein posttranslational modifications: the chemistry of proteome diversifications. Angew. Chem. Int. Ed. 2005, 44, 7342–7372.

2. Eberharter, A.; Becker, P. B., Histone acetylation: a switch between repressive and permissive chromatin. EMBO Rep. 2002, 3 (3), 224–229.

3. Hwang, C.-S.; Shemorry, A.; Varshavsky, A., N-terminal acetylation of cellular proteins creates specific degradation signals. Science 2010, 327, 973–977.

4. Ree, R.; Varland, S.; Arnesen, T., Spotlight on protein N-terminal acetylation. Exp. Mol. Med. 2018, 50 (7), 1–13.

5. Paik, W. K.; Paik, D. C.; Kim, S., Historical review: the field of protein methylation. Trends Biochem. Sci. 2007, 32 (3), 146–152.

6. Ohtsubo, K.; Marth, J. D., Glycosylation in cellular mechanisms of health and disease. Cell 2006, 126 (5), 855–867.

7. Humphrey, S. J.; James, D. E.; Mann, M., Protein phosphorylation: a major switch mechanism for metabolic regulation. Trends Endocrinol. Metab. 2015, 26 (12), 676–687.

8. Mann, M.; Jensen, O. N., Proteomic analysis of post-translational modifications. Nat. Biotech. 2003, 21, 255–261.

9. Kaji, A.; Kaji, H.; Novelli, G. D., A soluble amino acid incorporation system. Biochem. Biophys. Res. Com. 1963, 10, 406–409.

10. Kaji, H.; Novelli, G. D.; Kaji, A., A soluble amino acid-incorporating system from rat liver. Biochim. Biophys. Acta 1963, 76, 474–477.

11. Manahan, C. O.; App, A. A., An arginyl-transfer ribonucleic acid protein transferase from cereal embryos. Plant Physiol. 1973, 52 (1), 13–16.

12. Balzi, E.; Choder, M.; Chen, W. N.; Varshavsky, A.; Goffeau, A., Cloning and functional analysis of the arginyl-tRNA-protein transferase gene ATE1 of Saccharomyces cerevisiae. J. Biol. Chem. 1990, 265 (13), 7464–7471.

13. Van, V.; Smith, A. T., ATE1-Mediated Post-Translational Arginylation Is an Essential Regulator of Eukaryotic Cellular Homeostasis. ACS Chem. Biol. 2020, 15 (12), 3073–3085.

14. Bachmair, A.; Finley, D.; Varshavsky, A., In vivo half-life of a protein is a function of its amino-terminal residue. Science 1986, 234, 179–186.

15. Varshavsky, A., The N-end rule: functions, mysteries, uses. Proc. Natl. Acad. Sci. U.S.A. 1996, 93, 12142–12149.

16. Varshavsky, A., The N-end rule pathway and regulation by proteolysis. Protein Sci. 2011, 20 (8), 1298–1345.

17. Rao, H.; Uhlmann, F.; Nasymth, K.; Varshavsky, A., Degradation of a cohesin subunit by the N-end rule pathway is essential for chromosome stability. Nature 2001, 410, 955–960.

18. Kumar, A.; Birnbaum, M. D.; Patel, D. M.; Morgan, W. M.; Singh, J.; Barrientos, A.; Zhang, F., Posttranslational arginylation enzyme Ate1 affects DNA mutagenesis by regulating the stress response. Cell Death and Disease 2016, 7, e2378.

19. Kwon, Y. T.; Kashina, A. S.; Davydov, I. V.; Hu, R.-G.; An, J. Y.; Seo, J. W.; Du, F.; Varshavsky, A., An Essential Role of N-terminal Arginylation in Cardivascular Development. Science 2002, 297, 96–99.

20. Wang, J.; Pejaver, V. R.; Dann, G. P.; Wolf, M. Y.; Kellis, M.; Huang, Y.; Garcia, B. A.; Radivojac, P.; Kashina, A., Target site specificity and in vivo complexity of the mammalian arginylome. Sci. Rep. 2018, 8 (1), 16177.

21. Wang, J.; Han, X.; Leu, N. A.; Sterling, S.; Kurosaka, S.; Fina, M.; Lee, V. M.; Dong, D. W.; Yates, J. R.; Kashina, A., Protein arginylation targets alpha synuclein, facilitates normal brain health, and prevents neurodegeneration. Sci. Rep. 2017, 7 (1), 11323.

22. Bongiovanni, G.; Fidelio, G. D.; Barra, H. S.; Hallak, M. E., The post-translational incorporation of arginine into a beta-amyloid peptide increases the probability of alphahelix formation. Neuroreport 1995, 7 (1), 326–328.

23. Karakozova, M.; Kozak, M.; Wong, C. C. L.; Bailey, A. O.; Yates III, J. R.; Mogilner, A.; Zebroski, H.; Kashina, A., Arginylation of β-actin regulates actin cytoskeleton and cell motility. Science 2006, 313 (5784), 192–196.

24. Saha, S.; Mundia, M. M.; Zhang, F.; Demers, R. M.; Korobova, F.; Svitkina, T.; Perieteanu, A. A.; Dawson, J. F.; Kashina, A., Arginylation regulates intracellular actin polymer level by modulating actin properties and binding of capping and severing proteins. Mol. Biol. Cell 2010, 21 (8), 1350–1361.

25. Holman, T. J.; Jones, P. D.; Russell, L.; Medhurst, A.; Tomás, S. Ú.; Talloji, P.; Marquez, J.; Schmuths, H.; Tung, S.-A.; Taylor, I.; Footitt, S.; Bachmair, A.; Theodoulou, F. L.; Holdsworth, M. J., The N-end rule pathway promotes seed germination and establishment through removal of ABA sensitivity in Arabidopsis. Proc. Natl. Acad. Sci. U.S.A. 2009, 106 (11), 4549–4554.

26. White, M. D.; Klecker, M.; Hopkinson, R. J.; Weits, D. A.; Mueller, C.; Naumann, C.; O’Neill, R.; Wickens, J.; Yang, J.; Brooks-Bartlett, J. C.; Garman, E. F.; Grossman, T. N.; Dissmeyer, N.; Flashman, E., Plant cysteine oxidases are dioxygenases that directly enable arginyl transferase-catalysed arginylation of N-end rule targets. Nat. Commun. 2017, 8 (14690), doi: 10.1038/ncomms14690.

27. Holdsworth, M. J.; Gibbs, D. J., Comparative biology of oxygen sensing in plants and animals. Curr. Biol. 2020, 30 (8), R362–R369.

28. Berleth, E. S.; Li, J.; Braunscheidel, J. A.; Pickart, C. M., A reactive nucleophile proximal to vicinal thiols is an evolutionarily conserved feature in the mechanism of Arg aminoacyl-tRNA protein transferase. Arch. Biochem. Biophys. 1992, 298 (2), 498–504.

29. Li, J.; Pickart, C. M., Binding of phenylarsenoxide to Arg-tRNA protein transferase is independent of vicinal thiols. Biochemistry 1995, 34, 15829–15837.

30. Wang, J.; Han, X.; Wong Catherine C. L.; Cheng, H.; Aslanian, A.; Xu, T.; Leavis, P.; Roder, H.; Hedstrom, L.; Yates John R.; Kashina, A., Arginyltransferase ATE1 Catalyzes Midchain Arginylation of Proteins at Side Chain Carboxylates In Vivo. Chemistry & Biology 2014, 21 (3), 331–337.

31. Rai, R.; Mushegian, A.; Makarova, K.; Kashina, A., Molecular dissection of arginyltransferases guided by similarity to bacterial peptidoglycan synthases. EMBO Reports 2006, 7 (8), 800–805.

32. Van, V.; Brown, J. B.; Rosenbach, H.; Mohamed, I.; Ejimogu, N.-E.; Bui, T. S.; Szalai, V. A.; Chacón, K. N.; Span, I.; Smith, A. T., Iron sulfur clusters are involved in post-translational arginylation. bioRxiv 2021, DOI: 10.1101/2021.04.13.439645.

33. Varadi, M.; Anyango, S.; Deshpande, M.; Nair, S.; Natassia, C.; Yordanova, G.; Yuan, D.; Stroe, O.; Wood, G.; Laydon, A.; Ží;dek, A.; Green, T.; Tunyasuvunakool, K.; Petersen, S.; Jumper, J.; Clancy, E.; Green, R.; Vora, A.; Lutfi, M.; Figurnov, M.; Cowie,; Hobbs, N.; Kohli, P.; Kleywegt, G.; Birney, E.; Hassabis, D.; Velankar, S., AlphaFold Protein Structure Database: massively expanding the structural coverage of protein-sequence space with high-accuracy models. Nucl. Acids Res. 2022, 50 (D1), D439–D444.

34. Ud-Din, A. I. M. S.; Tikhomirova, A.; Roujeinkova, A., Structure and functional diversity of GCN5-related N-acetyltransferases (GNAT). Int. J. Mol. Sci. 2016, 17, 1018.

35. Watanabe, K.; Toh, Y.; Suto, K.; Shimizu, Y.; Oka, N.; Wada, T.; Tomita, K., Protein-based peptide-bond formation by aminoacyl-tRNA protein transferase. Nature 2007, 449, 867–872.

36. Holm, L.; Rosenstrom, P., Dali server: conservation mapping in 3D. Nucl. Acids Res. 2010, 38, W545–W549.

37. Saha, S.; Wang, J.; Buckley, B.; Wang, Q.; Lilly, B.; Chernov, M.; Kashina, A., Small molecule inhibitors of arginyltransferase regulate arginylation-dependent protein degradation, cell motility, and angiogenesis. Biochem. Pharamacol. 2012, 83, 866–873.

38. Avcilar-Kucukgoze, I.; Gamper, H.; Polte, C.; Ignatova, Z.; Kraetzner, R.; Shtutman, M.; Hou, Y.-M.; Dong, D. W.; Kashina, A., tRNAArg-Derived Fragments Can Serve as Arginine Donors for Protein Arginylation. Cell Chem. Biol. 2020, 27, 1–11.

39. Fung, A. W. S.; Leung, C. C. Y.; Fahlman, R. P., The determination of tRNALeu recognition nucleotides for Escherichia coli L/F transferase. RNA 2014, 20 (8), 1210–1222.

40. Varshavsky, A., N-degron and C-degron pathways of protein degradation. Proc. Natl. Acad. Sci. U.S.A. 2019, 116 (2), 358–366.

41. An, J. Y.; Seo, J. W.; Tasaki, T.; Lee, M. J.; Varshavsky, A.; Kwon, Y. T., Impaired neurogenesis and cardiovascular development in mice lacking the E3 ubiquitin ligases UBR1 and UBR2 of the N-end rule pathway. Proc. Natl. Acad. Sci. U.S.A. 2006, 103, 6212–6217.

42. Park, S.-E.; Kim, J.-M.; Seok, O.-H.; Cho, H.; Wadas, B.; Kim, S.-Y.; Varshavsky, A.; Hwang, C.-S., Control of mammalian G protein signaling by N-terminal acetylation and the N-end rule pathway. Science 2015, 347 (6227), 1249–1252.

43. Fina, M. E.; Wang, J.; Nikonov, S. S.; Sterling, S.; Vardi, N.; Kashina, A.; Dong, D. W., Arginyltransferase (Ate1) regulates the RGS7 protein level and the sensitivity of light-evoked ON-bipolar responses. Sci. Rep. 2021, 11 (1), 9376.

44. Wong, C. C. L.; Xu, T.; Rai, R.; Bailey, A. O.; Yates, J. R.; Wolf, Y. I.; Zebroski, H.; Kashina, A., Global Analysis of Posttranslational Protein Arginylation. PLoS Biology 2007, 5 (10), e258.

45. Rai, R.; Zhang, F.; Colavita, K.; Leu, N. A.; Kurosaka, S.; Kumar, A.; Birnbaum, M. D.; Győrffy, B.; Dong, D. W.; Shtutman, M.; Kashina, A., Arginyltransferase suppresses cell tumorigenic potential and inversely correlates with metastases in human cancers. Oncogene 2016, 35 (31), 4058–4068.

46. Kishimoto, N.; Okano, R.; Akita, A.; Miura, S.; Irie, A.; Takamune, N.; Misumi, S., Arginyl-tRNA-protein transferase 1 contributes to governing optimal stability of the human immunodeficiency virus type 1 core. Retrovirology 2021, 18, doi: 10.1186/s12977-021-00574-0.

47. Vedula, P.; Tang, H.-Y.; Speicher, D. W.; Kashina, A., Protein posttranslational signatures identified in COVID-19 patient plasma. Front. Cell. Dev. Biol. 2022, DOI: 10.3389/fcell.2022.807149.

48. Brower, C. S.; Rosen, C. E.; Jones, R. H.; Wadas, B. C.; Piatkov, K. I.; Varshavsky, A., Liat1, an arginyltransferase-binding protein whose evolution among primates involved changes in the numbers of its 10-residue repeats. Proc. Natl. Acad. Sci. U.S.A. 2014, 111 (46), E4936–E4945.

49. Linkous, R. O.; Sestok, A. E.; Smith, A. T., The crystal structure of Klebsiella pneumoniae FeoA reveals a site for proteinLJprotein interactions. Proteins 2019, 87 (11), 897–903.

50. Winter, G., Xia2: an expert system for macromolecular crystallography data reduction.. J. Appl. Cryst. 2010, 43, 186–190.

51. Adams, P. D.; Afonine, P. V.; Bunkóczi, G.; Chen, V. B.; Davis, I. W.; Echols, N.; Headd, J. J.; Hung, L.-W.; Kapral, G. J.; Grosse-Kunstleve, R. W.; McCoy, A. J.; Moriarty, N. W.; Oeffner, R.; Read, R. J.; Richardson, D. C.; Richardson, J. S.; Terwilliger, T. C.; Zwart, P. H., PHENIX: a comprehensive Python-based system for macromolecular structure solution. Acta. Cryst. D. 2010, 66, 213–221.

52. Emsley, P.; Cowtan, K., Coot: model-building tools for molecular graphics. Acta. Cryst. D. 2004, 60, 2126–2132.

53. Pettersen, E. F.; Goddard, T. D.; Huang, C. C.; Couch, G. S.; Greenblatt, D. M.; Meng, E. C.; Ferrin, T. E., UCSF Chimera--a visualization system for exploratory research and analysis. J. Comput. Chem. 2004, 25 (13), 1605–1612.

54. Stivala, A.; Wybrow, M.; Wirth, A.; Whisstock, J.; Stuckey, P., Automatic generation of protein structure cartoons with Pro-origami. Bioinformatics 2011, 27 (23), 3315–3316.

55. Förster, S.; Timmann, A.; Konrad, M.; Schellbach, C.; Meyer, A.; Funari, S. S.; Mulvaney, P.; Knott, R., Scattering curves of ordered mesoscopic materials. J. Phys. Chem. B 2005, 109 (4), 1347–60.

56. Förster, S.; Apostol, L.; Bras, W., Scatter: software for the analysis of nano- and mesoscale small-angle scattering. J. Appl. Crystallogr. 2010, 43 (3), 639–646.

57. Svergun, D. I., Determination of the regularization parameter in indirect-transform methods using perceptual criteria. J. Appl. Crystallogr. 1992, 25 (4), 495–503.

58. Svergun, D. I.; Petoukhov, M. V.; Koch, M. H., Determination of domain structure of proteins from X-ray solution scattering. Biophys. J. 2001, 80 (6), 2946–53.

59. Volkov, V. V.; Svergun, D. I., Uniqueness of ab initio shape determination in small-angle scattering. J. Appl. Crystallogr. 2003, 36 (3 Part 1), 860–864.

60. Kozin, M. B.; Svergun, D. I., Automated matching of high- and low-resolution structural models. J. Appl. Crystallogr. 2001, 34 (1), 33–41.

61. Waterhouse, A. M.; Procter, J. B.; Martin, D. M. A.; Clamp, M.; Barton, G. J., Jalview Version 2 - a multiple sequence alignment editor and analysis workbench. Bioinformatics 2009, 25 (9), 1189–1191.

62. Larkin, M. A.; Blackshields, G.; Brown, N. P.; Chenna, R.; McGettigan, P. A.; McWilliam, H.; Valentin, F.; Wallace, I. M.; Wilm, A.; Lopez, R.; Thompson, J. D.; Gibson, T. J.; Higgins, D. G., Clustal W and Clustal X version 2.0. Bioinformatics 2007, 23 (21), 2947–2948.

63. Henikoff, S.; Henikoff, J. G., Amino acid substitution matrices from protein blocks. Proc. Natl. Acad. Sci. USA 1992, 89, 10915–10919.

64. Crooks, G. E.; Hon, G.; Chandonia, J. M.; Brenner, S. E., WebLogo: A sequence logo generator. Genome Res. 2004, 14, 1188–1190.

65. Schneider, T. D.; Stephens, R. M., Sequence logos: A new way to display consensus sequences. Nucleic Acids Res. 1990, 18 (20), 6097–6100.

