## Supplemental Information for "The structure of arginyltransferase 1 (ATE1)"

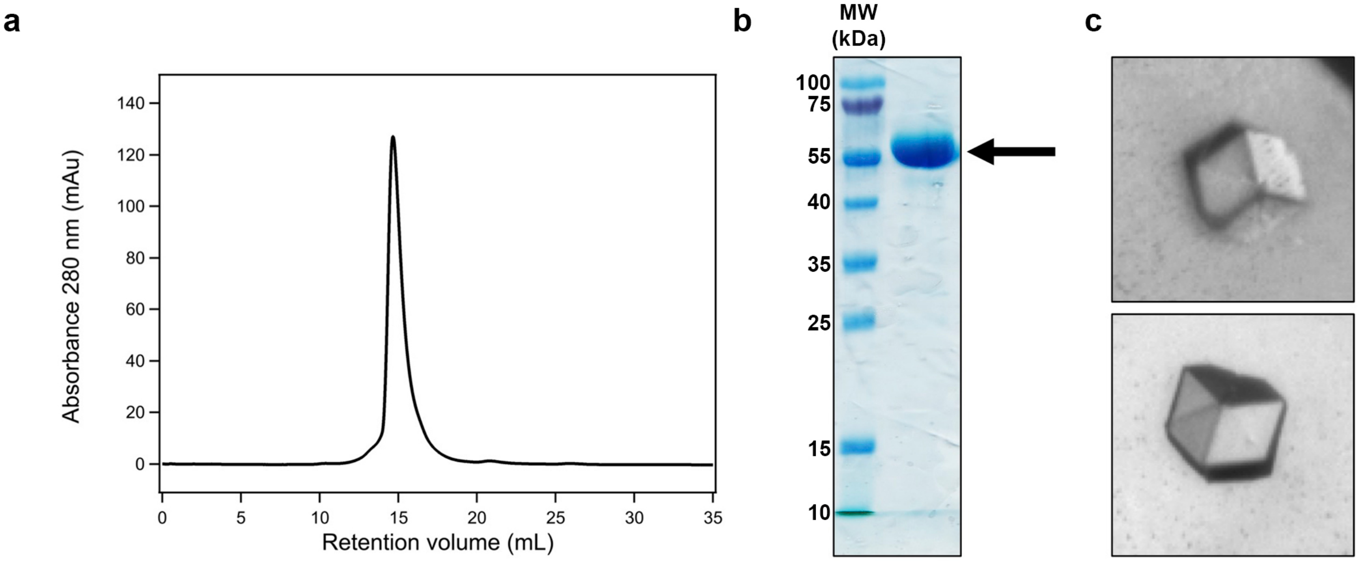


**Supplementary Figure 1**. Heterologously-produced and purified *Sc*ATE1 is highly pure, monodisperse, and crystallizes reproducibly. **a**. Size-exclusion chromatogram of purified *Sc*ATE1. The retention volume observed is consistent with monomeric *Sc*ATE1 (≈ 60 kDa). **b**. 15% SDS-PAGE analysis of the final, SEC-purified *Sc*ATE1. The single, major band indicating the presence of highly-pure *Sc*ATE1 is indicated by an arrow. **c**. Examples of *Sc*ATE1 crystals produced from the purified protein analyzed in panels **a** and **b**.

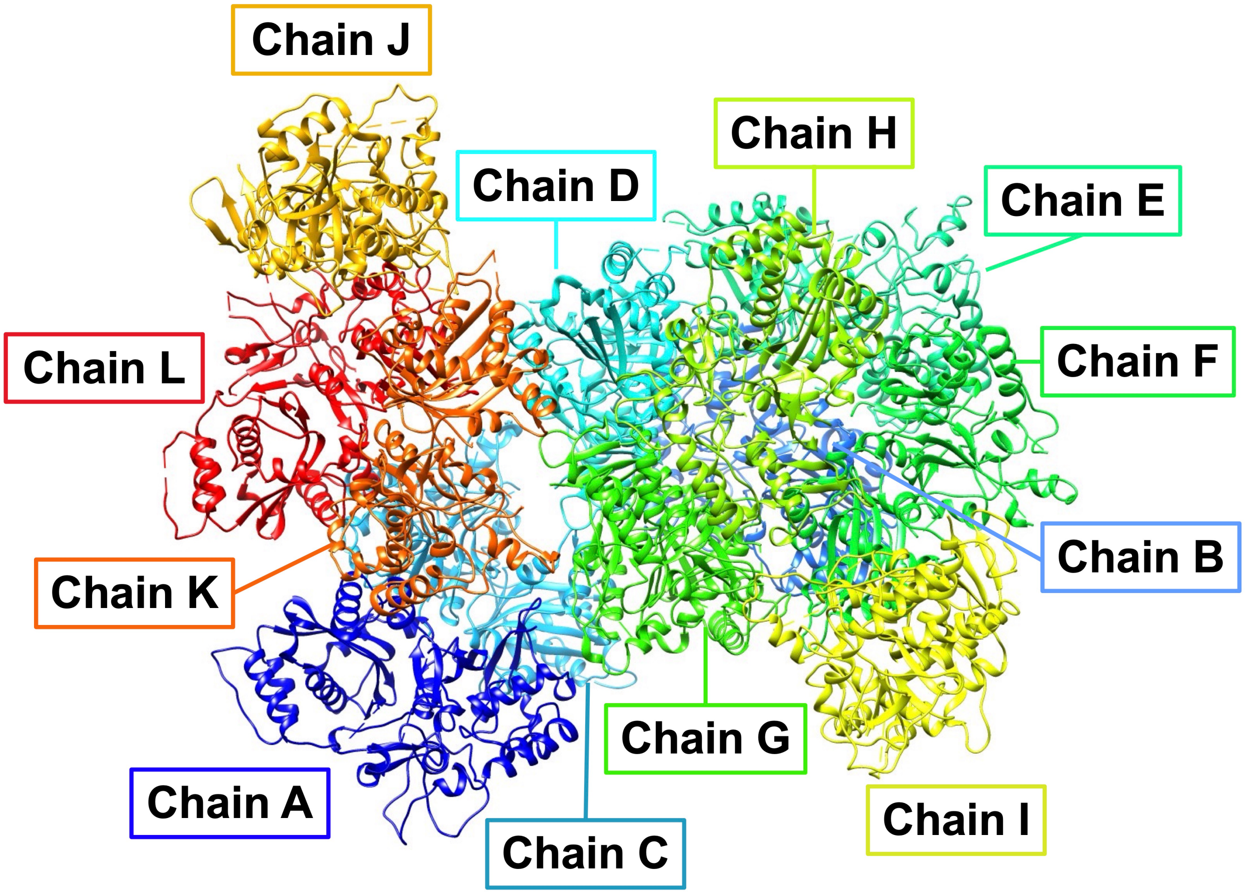


**Supplementary Figure 2**. The asymmetric unit (ASU) of *Sc*ATE1 crystallized in the $P4_3$ space group. The *Sc*ATE1 ASU is composed of 12 *Sc*ATE1 polypeptides (labeled as chains A-L). Each chain is individually colored from dark blue (chain A) to red (chain L). Regions of disorder that are unmodeled in the final structure are indicated by dashed lines.

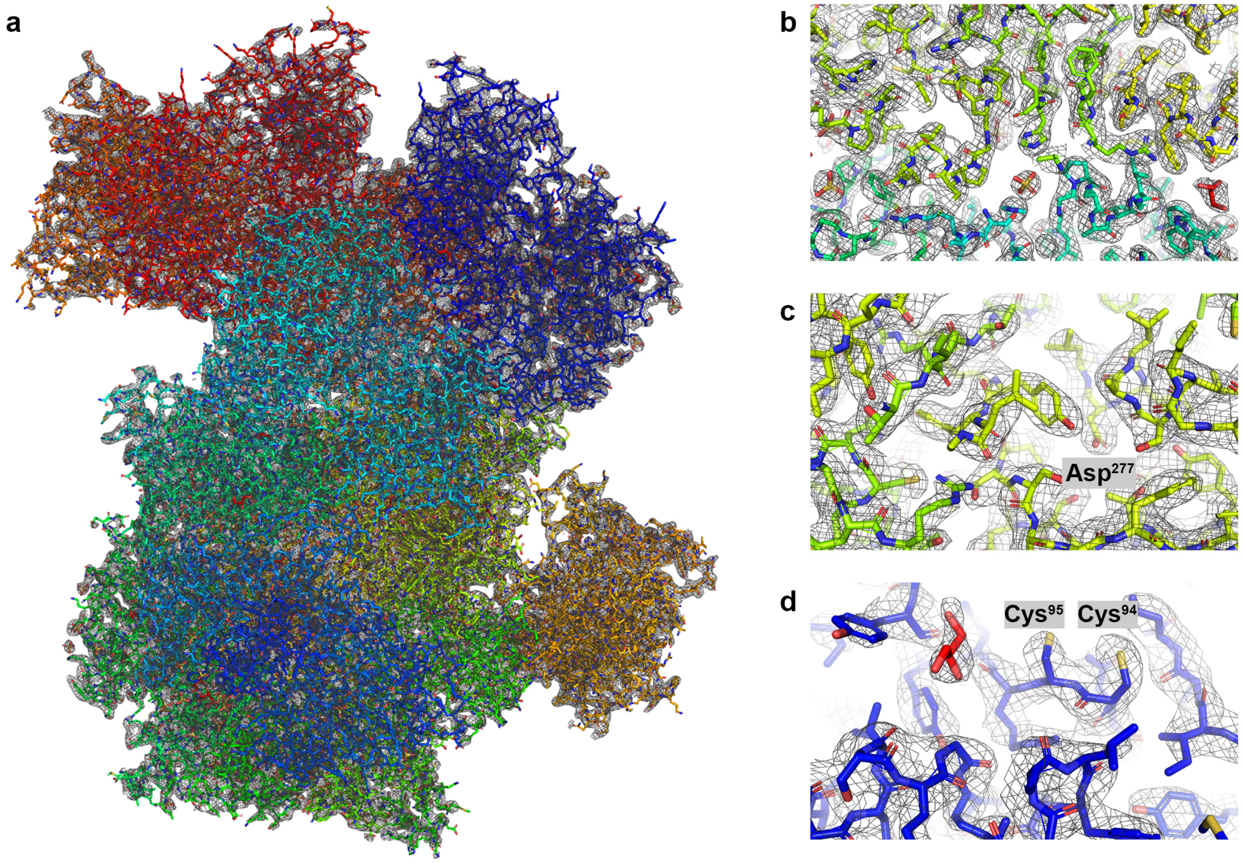


**Supplementary Figure 3**. The electron density map of *Sc*ATE1. The 2$F_o$-$F_c$ electron density map (gray) is contoured at 1σ, and water, sulfate, and glycerol molecules are all explicitly shown. **a**. A single asymmetric unit (12 polypeptides) of *Sc*ATE1, color-coded differentially based on polypeptide chain. **b**. Example of the 2$F_o$-$F_c$ electron density map at the interface of 4 chains of *Sc*ATE1. **c**. Example of the 2$F_o$-$F_c$ electron density map at a key active site residue (Asp$^{277}$) and its surrounding hydrophilic pocket. **d**. Example of the 2$F_o$-$F_c$ electron density map at the vicinal thiols (Cys$^{94}$ and Cys$^{95}$), which are part of the binding site of the [Fe-S] cluster. This structure is apo (lacks the [Fe-S] cluster), as evidenced by the absence of electron density for the cluster and the absence of electron density of the dynamic/disordered Cys$^{20}$xxCys$^{23}$ motif in every chain of the structure.

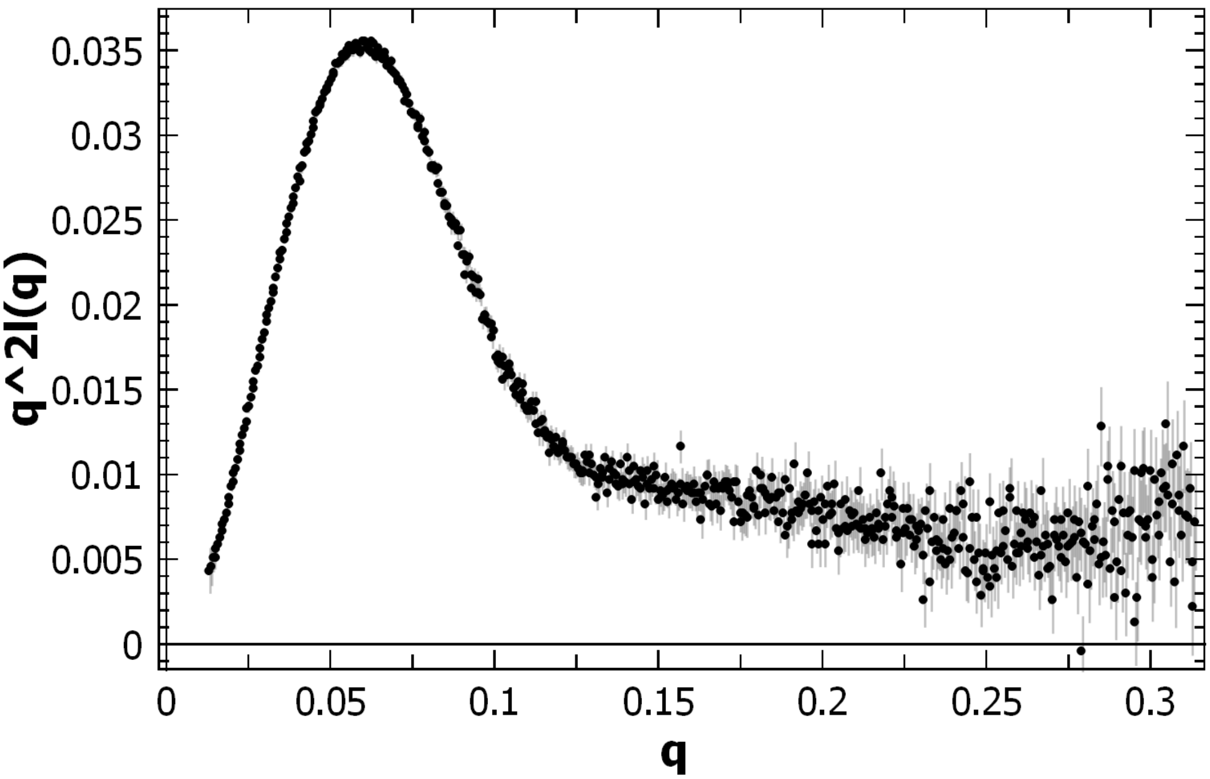


**Supplementary Figure 4**. The Kratky plot of *Sc*ATE1. The plot is Gaussian in shape suggesting a folded, globular protein in solution, and there is no indication of the presence of additional, distinct domains.

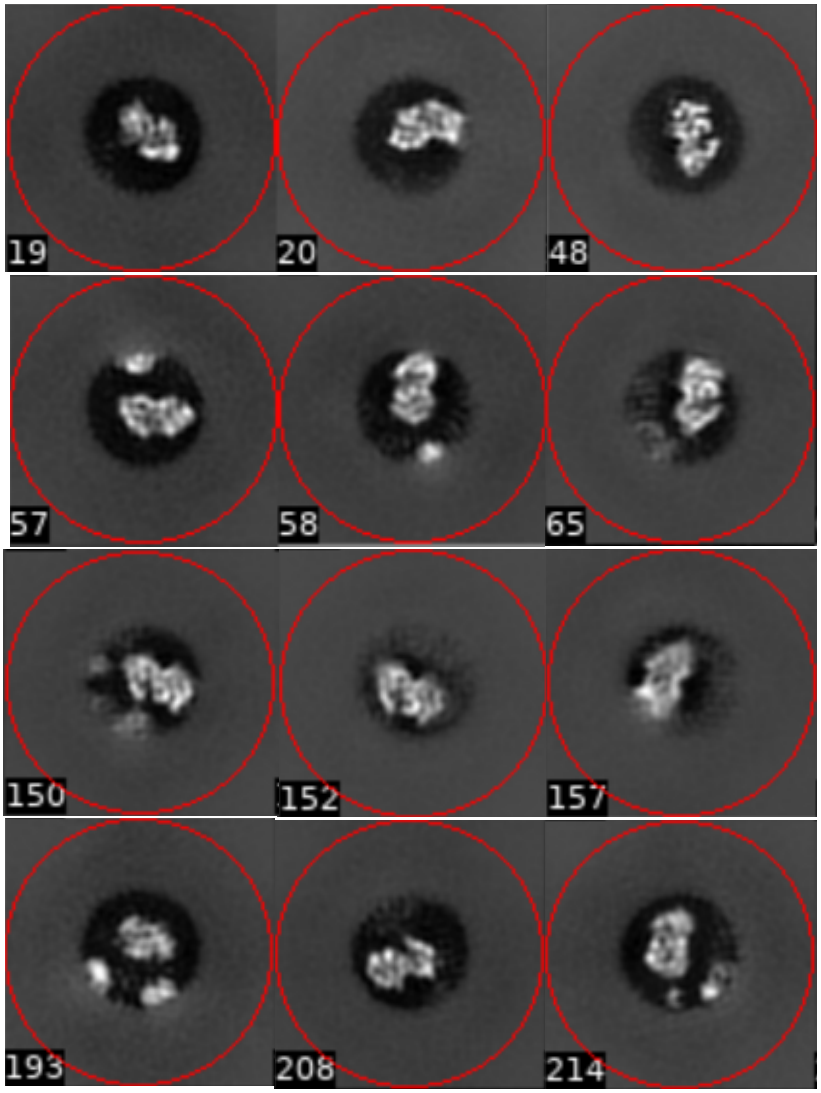


**Supplementary Figure 5**. Select cryo-EM 2D class average images of *Sc*ATE1. Particle images display a bilobed, roughly globular shape in the center of a masking radius (black circle) of 60 Å.

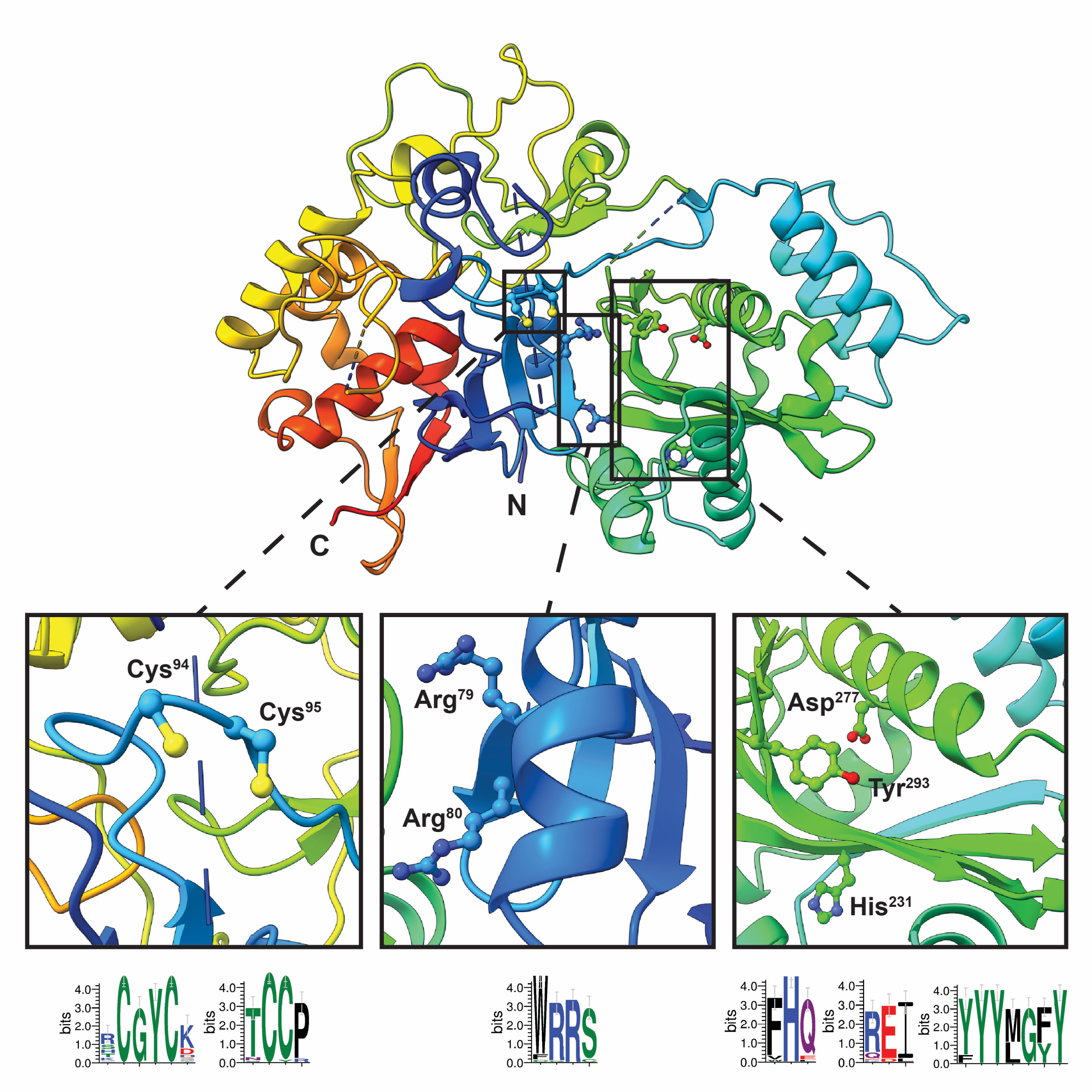


**Supplementary Figure 6.** Residues of interest in the *Sc*ATE1 polypeptide. Top panel shows the full X-ray crystal structure of a single *Sc*ATE1 polypeptide colored in rainbow, with regions of interest in black outlined boxes. Bottom left panel: close-up view of $Cys^{94}$ and $Cys^{95}$, with the Weblogo highlighting sequence conservation of both Cys motifs known to coordinate the [Fe-S] cluster below. Bottom middle panel: close-up view of $Arg^{79}$ and $Arg^{80}$, the latter of which is believed to be the residue responsible for recognition of the negatively charged N-terminal substrates. The conservation of both residues is highlighted in Weblogo below. Bottom right panel: close-up view of the catalytic site of the *Sc*ATE1 GNAT fold highlighting $Asp^{277}$, $Tyr^{293}$, and $His^{231}$. with the Weblogo below.

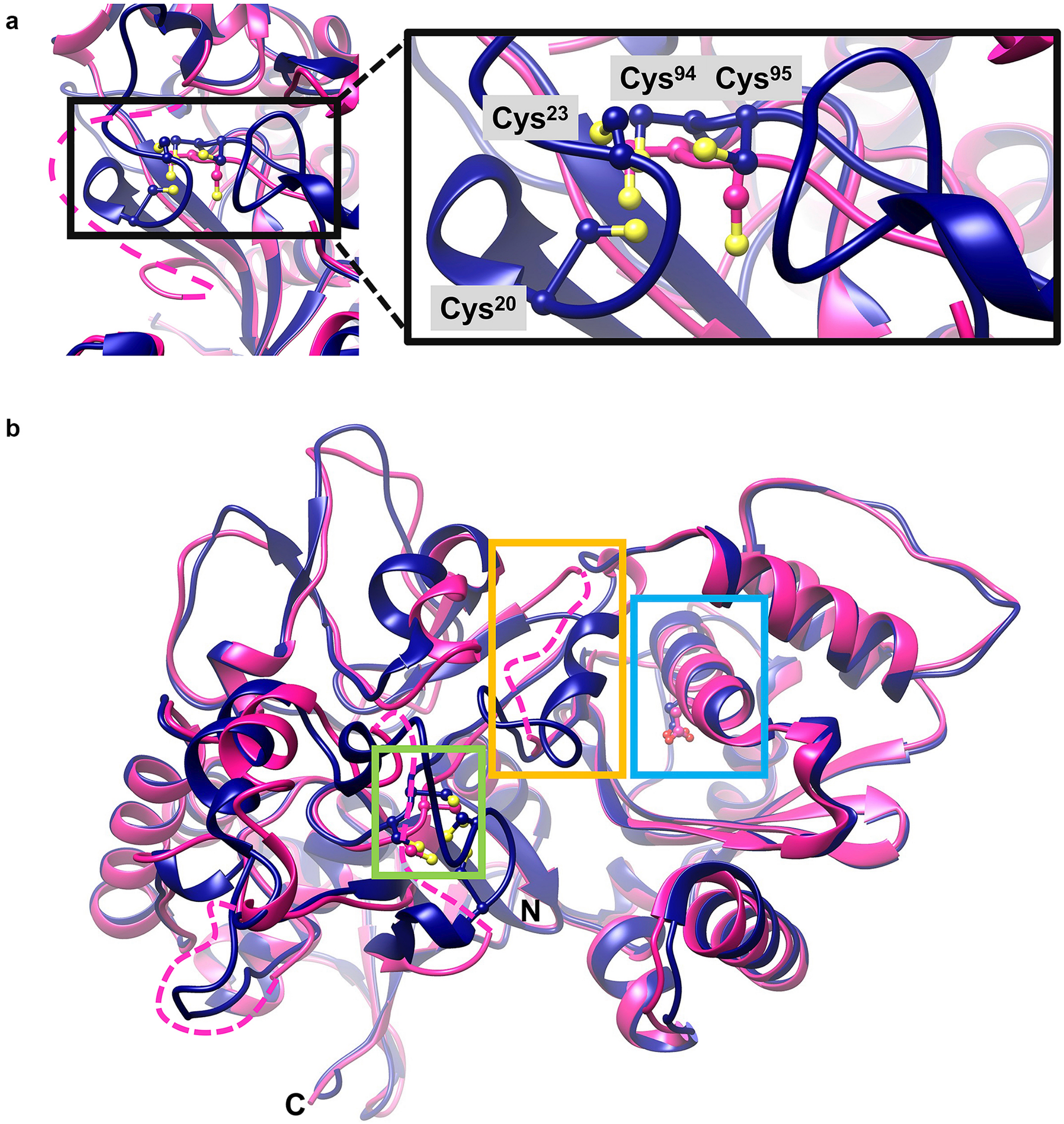


**Supplementary Figure 7**. Comparison of the X-ray crystal structure and the AlphaFold model of *Sc*ATE1. The X-ray structure is colored magenta and the AlphaFold model is colored blue. **a**. Close-up of the N-terminal regulatory domain of *Sc*ATE1 that binds an [Fe-S] cluster. The expanded region highlights the location of four Cys residues necessary for cluster binding: $Cys^{20}$, $Cys^{23}$, $Cys^{94}$, and $Cys^{95}$. In the crystal structure, the loop containing $Cys^{20}$ and $Cys^{23}$ is disordered (represented by a magenta dashed line), but the loop is modeled in the AlphaFold model, strongly suggesting co-localization of these residues for cluster binding. **b**. We hypothesize that the binding of the [Fe-S] cluster in the N-terminal regulatory domain (green box) is communicated through a second, disordered region (yellow box) that is located between the cluster-binding site and the active site GNAT fold containing the key $Asp^{277}$ residue (blue box).

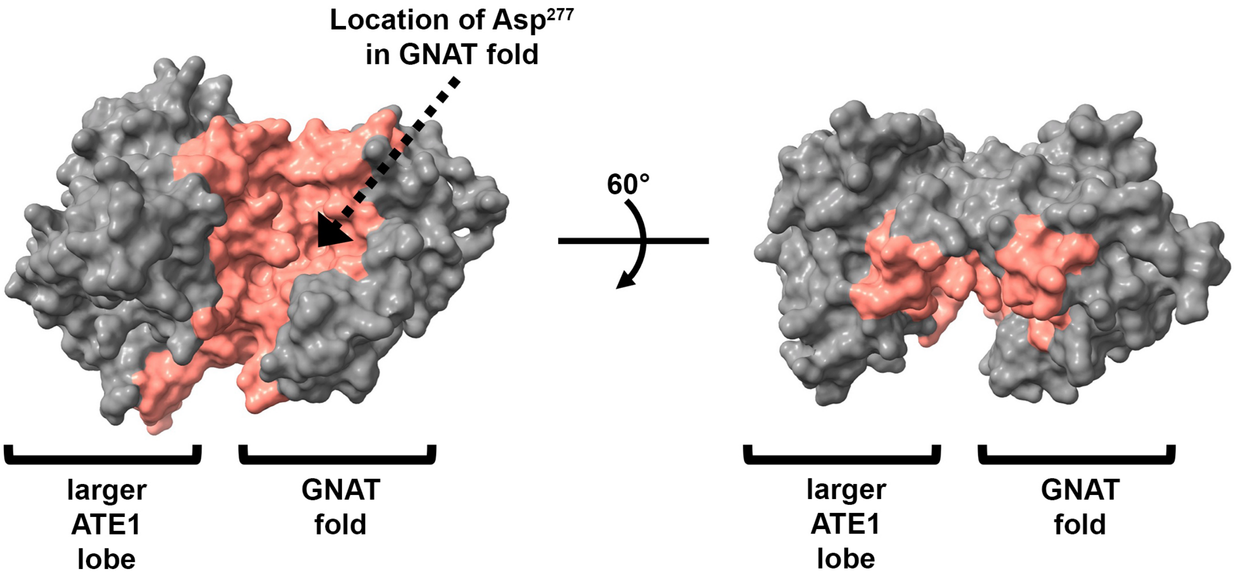


**Supplementary Figure 8**. The Van der Waals surface representation of *Sc*ATE1 and the large cleft between the two lobes of the protein. The larger ATE1 lobe comprises the N-terminal regulatory domain composed of residues 1-103 and the C-terminal domain of unknown function composed of residues 307-502. The GNAT fold comprises residues 107-307. The sizable cleft between the two lobes is colored in salmon and is estimated at 6700 Å$^3$ in volume. The right panel represents a 60° rotation about the X axis of the figure in the left panel.

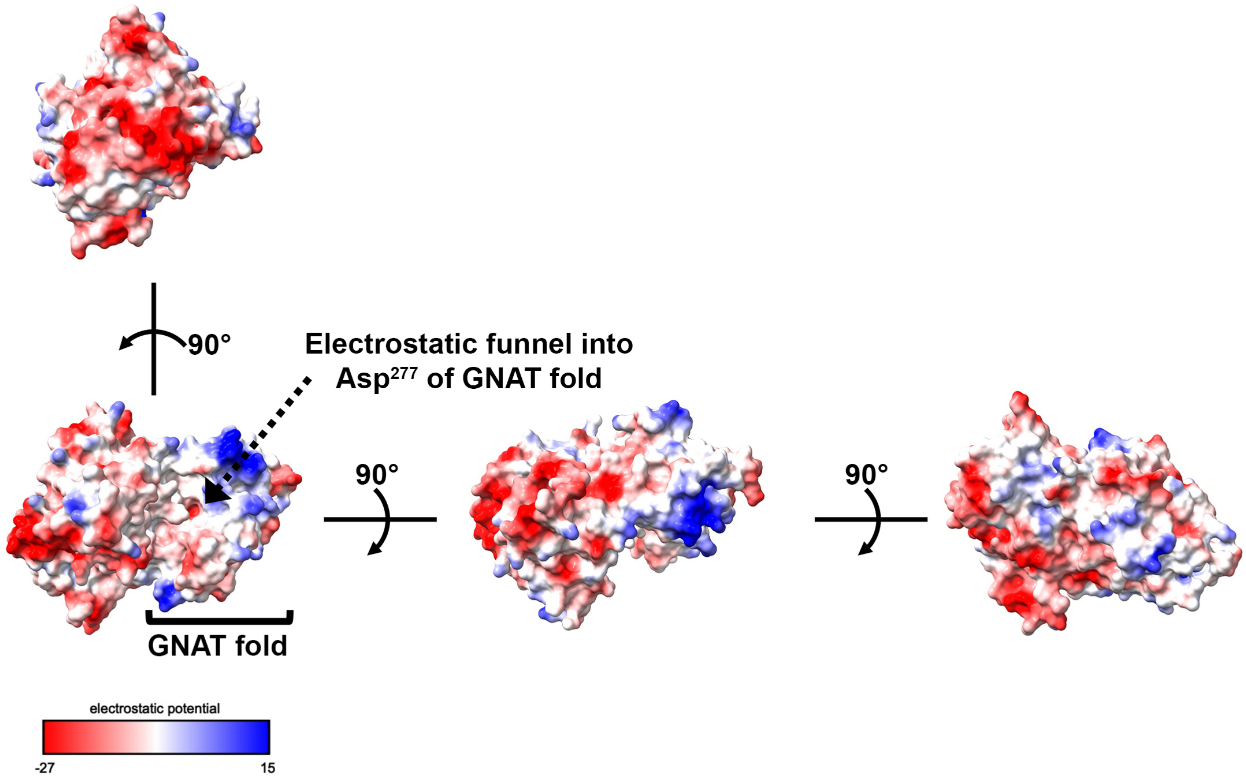


**Supplementary Figure 9**. The electrostatic surface of the *Sc*ATE1 X-ray crystal structure reveals a putative location for tRNA binding. Lower left panel: the Van der Waals surface representation of *Sc*ATE1 is color-coded based on electrostatic potential from -27 kcal/(mol·e$^-$) (red) to +15 kcal/(mol·e$^-$) (blue). The GNAT fold of *Sc*ATE1 is bracketed, and an electrostatic funnel into Asp$^{277}$ of the GNAT fold is indicated. The middle panel represents a 90° rotation about the X axis of the lower left panel. The right panel represents a 90° rotation about the X axis of the middle panel. The upper left panel represents a 90° rotation about the Y axis of the lower left panel. The majority of the *Sc*ATE1 surface is strongly electronegative (red), precluding these portions of the surface from binding the negatively-charged tRNA. The exception to this observation is an electropositive patch along the GNAT that funnels into Asp$^{277}$, the putative binding site of the 3' aminoacylated end of the tRNA.

| | ***Sc*ATE1** |
|---|---|
| **Data collection** | |
| Wavelength (Å) | 0.97872 |
| Space group | $P4_3$ |
| Cell dimensions | |
| *a*, *b*, *c* (Å) | 235.27, 235.27, 171.10 |
| $\alpha$, $\beta$, $\gamma$ (°) | 90, 90, 90 |
| $R_{merge}$ | 0.167 (2.288) |
| $R_{pim}$ | 0.065 (0.879) |
| I / σ (I) | 11.0 (1.0) |
| Completeness (%) | 100.0 (100.0) |
| Multiplicity | 7.7 (7.8) |
| CC½ (%) | 99.7 (33.0) |
| **Refinement** | |
| Resolution (Å) | 68.23 – 2.85 (2.90 – 2.85) |
| No. unique reflections | 216,770 (21,567) |
| $R_{work}$ | 0.199 (0.319) |
| $R_{free}$ | 0.257 (0.367) |
| Macromolecules per ASU | 12 |
| No. non-H atoms/molecules | |
| Protein atoms | 44714 |
| Water molecules | 64 |
| Sulfate molecules | 203 |
| Glycerol molecules | 41 |
| Average *B*-factors ($Å^2$) | |
| Protein | 88.50 |
| Water | 59.91 |
| Sulfate and glycerol | 147.78 |
| R.m.s. deviations | |
| Bond lengths (Å) | 0.010 |
| Bond angles (°) | 1.230 |
| Ramachandran plot (%) | 93.0 / 6.9 / 0.1 |

**Supplementary Table 1**. Data collection and refinement statistics of apo *Sc*ATE1. Values in parentheses are for the highest resolution shell.
